# Time-limited alterations in cortical activity of a Knock-in mice model of *KCNQ2-*related Developmental and Epileptic Encephalopathy

**DOI:** 10.1101/2020.05.12.090464

**Authors:** Najoua Biba, Hélène Becq, Marie Kurz, Emilie Pallesi, Laurent Villard, Mathieu Milh, Pierre-Pascal Lenck Santini, Laurent Aniksztejn

**Affiliations:** INSERM, INMED, Aix-Marseille University, Marseille, France; Aix-Marseille Univ, INSERM, MMG, Marseille, France; Department of Medical Genetics, La Timone Childrens’s Hospital, Marseille, France; Department of Pediatric Neurology, La Timone Children’s Hospital, Marseille, France

## Abstract

*De novo* missense variants in the *KCNQ2* gene encoding the Kv7.2 subunit of the voltage-gated potassium Kv7/M channel are the main cause of Developmental and Epileptic Encephalopathy (DEE). *KCNQ2* related-DEE is characterized by pharmaco-resistant neonatal seizures associated with a developmental delay. While seizures usually resolve some weeks or months after birth, cognitive/behavioral deficits persist. To better understand the cellular mechanisms underlying *KCNQ2-associated* network dysfunction and their progression over time, we investigated *in vivo,* using local field potential recordings of freely moving animals, and *ex-vivo* in layers II/III and V of motor cortical slices, using patch-clamp recordings, the electrophysiological properties of pyramidal cells from a heterozygous knock-in (KI) mouse model carrying the p.T274M pathogenic variant during neonatal, post-weaning and juvenile developmental stages. We found that KI mice displayed spontaneous seizures preferentially at post-weaning rather than at juvenile stages. At the cellular level, the variant led to a reduction in M current density/chord conductance and to an increase in neuronal excitability. These alterations were observed already during the neonatal period in pyramidal cells of layers II / III and during post-weaning stage in pyramidal cells of layer V. Moreover there was an increase in the frequency of spontaneous network driven events mediated by GABA receptors in the layers II/III suggesting that the excitability of some interneurons was also increased. However, all these alterations were time limited and no more observed in layers II/III and V of juvenile mice. At this stage, M-current density and neuronal excitability were not different from the measurements made in juvenile wild-type mice. Thus our data indicate that the action of the variant on neuronal activity is developmentally regulated and that some cellular mechanisms leading to the recovery of Kv7/M channels function took place during brain maturation of KI mice. These results raise the possibility that the age related seizure remission observed in *KCNQ2*-related DEE patient results also from a time limited alteration of Kv7 channels activity and neuronal excitability.

## Introduction

Channelopathies represent one of the major causes of neurological disorders including Developmental and Epileptic Encephalopathy (DEE), a group of severe and intractable diseases that associate severe epilepsy with an alteration of cognitive/sensory and motor functions. Among dozens of genes encoding for ion channels, *KCNQ2* is probably the most frequently one associated to DEE with neonatal onset (Weckhuysen et al., 2012; Kato et al., 2013; Milh et al., 2013; Allen et al., 2014, 2020; McTague et al., 2016; Dirkx et al., 2020; see also http://www.rikee.org/). *KCNQ2-*related DEE is characterized by seizures starting during the first days of life and often a highly abnormal EEG showing a suppression-burst pattern. As patients age, there is usually a positive evolution in terms of seizures and EEG abnormalities. However despite seizure resolution, the neurological outcome remains poor, with severe motor and intellectual disabilities (Dirkx et al., 2021, Boets et al., 2021).

*KCNQ2* is one of the four genes identified in the central nervous system with *KCNQ3-5*, each gene encoding respectively the Kv7.2-7.5 subunits of the Kv7 channels (Jentsch, 2000). These subunits are abundant at axon initial segment (AIS), nodes of Ranvier, and synaptic terminals ( Devaux et al. 2004; Battefeld et al., 2014; Shah et al., 2008; Huang and Trussell, 2011; Martinello et al., 2019).There are less expressed at the soma but they are absent in dendrites of cortical pyramidal cells while in cortical interneurons these channels are detected in these subcellular regions (Lawrence et al., 2006).

Functional Kv7 channels are composed of homomeric or heteromeric assemblies of 4 subunits including Kv7.2/Kv7.3 in many cortical neurons giving rise to the M current (Wang et al. 1998; Brown and Passmore, 2009; Battefeld et al., 2014; Soh et al., 2014). M current is a slowly activating and non-inactivating potassium current activating at subthreshold range of membrane potentials. Kv7/M channels control different aspects of neuronal excitability (Adams et al., 1982; Storm 1990; Jentsch, 2000; Greene and Hoshi 2017) including the resting membrane potential at the AIS and terminals as well as the spike threshold (Yue and Yaari, 2006; Shah et al., 2008; Hu and Bean 2018, Huang and Trussell, 2011; Martinello et al., 2019). At nodes of Ranvier, Kv7 channels increase the availability of Na^+^ channels and thus the amplitude of the propagating action potential (Battefeld et al., 2014). At synaptic terminals they can also control transmitter release (Vervaeke et al., 2006; Huang and Trussell, 2011; Martinello et al., 2019). Kv7 channels also reduce spike afterdepolarization, and mediate in some cells the medium afterhyperpolarization (mAHP) and contribute to the slow AHP (sAHP) (Storm, 1990; Gu et al., 2005; Yue and Yaari 2006, Kim et al., 2012). Thus, Kv7 channels serve as a brake for neuronal firing. These channels also influence other aspects of neuronal function such as the integration of excitatory post-synaptic potential in CA1 pyramidal cells (Hu et al., 2007; Shah et al., 2011) or the slow substhreshold resonance of cortical neurons at theta frequency (Hu et al., 2002).

Although the functional role of Kv7 channels and their different subunits is now well documented (Brown and Passmore, 2009; Peters et al. 2015; Soh et al.,2014, 2018; Fidzinski et al., 2015; Greene and Hoshi, 2017; Hou et al., 2021), the relationship between Kv7.2 variants and DEE remains poorly understood. Functional consequences of variants in the *KCNQ2* gene associated to DEE have widely been analyzed in heterologous cells and more recently in patient iPSC derived neurons (Dirkx et al., 2020; Simkin et al., 2021). In heterologous cells, these studies have shown that most of the variants exerted a loss of function effect reducing M current carried by heteromeric channels from a moderate level (∼25 %) to a strong level (>50%) (Miceli et al. 2013; Ohran et al., 2014; Abidi et al., 2015; Allen et al., 2020; Simkin et al., 2021). This is notably the case of the p.T274 M variant (c.821C>T in the *KCNQ2* gene). This variant exerts a dominant negative effect on wild type subunit reducing current amplitude by 60% in a heterozygous configuration that mimics patient’s situation and without any effects on the conductance-voltage relationship, protein production and membrane expression (Orhan et al., 2014). Recently, a knock-in mouse carrying the heterozygous p.T274M variant has been generated, the phenotype of which was reminiscent of some characteristics observed in *KCNQ2*-related DEE patients (Milh et al., 2020). In particular, spontaneous seizures have been recorded in juvenile KI mice and cognitive impairment observed in adults. Thus, this knock-in mouse represent a good model to better understand how a Kv7.2 pathogenic variant alters cortical network functioning and ultimately help at designing therapeutic options.

Here, we investigated both *in vivo* and *ex-vivo* in motor cortical slices the electrophysiological consequences of the p.T274M pathogenic variant on cortical activity. More precisely we analyzed the possible alterations that the variant may produce on pyramidal cells excitability, M current properties and spontaneous synaptic activities and performed a developmental study to see how these alterations, if any, evolved as animal age from neonatal to juvenile stages. In addition, given that expression of ion channels may not be equivalent in pyramidal cells of the different cortical layers we wondered if the effect of the variant is the same or differed between pyramidal cells of the layers II/III and of the layer V of the developing motor cortex. Finally since in patients, the frequency of seizures declined or remitted during childhood (Boets et al., 2021), we wondered if the occurrence of seizures is also developmentally regulated in these KI mice.

## Results

Pyramidal cells located in layers II/III and V of the developing motor cortex were recorded using patch-clamp technique in slices from mice aged one week (neonatal, postnatal day 7-9, PND7-9), three weeks (post-weaning, PND 19-21) and 4-5 weeks (juvenile, PND28-35). The properties of the cells obtained from wild type mice (*KCNQ2^WT/WT^*, hereinafter referred to as “wild type cells”) and from heterozygous knock-in *KCNQ2* mutant mice harboring the p.T274M variant (*KCNQ2^WT/T274M^*, hereinafter referred to as “mutant cells”) were compared. Furthermore, since the p.T274M variant exerts a loss of function effect on Kv7/M channels activity in heterologous systems (Orhan et al., 2014), we also analyzed the action of the potent Kv7/M channels blocker XE-991 (20 µM) on pyramidal cells properties of wild type mice to see if the variant and the blocker had similar electrophysiological consequences. For this purpose, slices obtained from *KCNQ2^WT/WT^* mice were superfused with the blocker (*KCNQ2^WT/WT^* in XE-991, hereinafter referred to as “XE-991 treated wild-type cells”).

### The p.T274M variant transiently increases the excitability of pyramidal cells of the layers II/III

At PND7-9, in cell-attached configuration, pyramidal cells of the layers II/III recorded in the three groups of slices were silent, there was not any spontaneous discharge in wild type cells recorded either in absence or in presence of XE-991, and in mutant cells (n = 17 wild-type cells/ 2 animals; n = 11 XE-991 treated wild-type cells/ 2 animals; n =17 mutant cells/3 animals; Fig 1Aa) indicating that the variant did not excite pyramidal cells at their resting membrane potential. In whole-cell configuration, compared to wild-type cells, mutant cells and XE-991 treated wild type cells exhibited a higher input resistance (Rm), a lower current threshold (rheobase) to elicit a single action potential (AP) by short 10 msec depolarizing current steps and a more hyperpolarized action potential membrane threshold (e.g., “Table 1”). We then analyzed the neuronal discharge elicited by the injection of depolarizing current steps command from 20 to 300 pA during 1 sec. We quantified the number of AP, the frequency of the discharge during the first 200 msec of the steps (initial frequency) and during the last 200 msec of the steps (final frequency). The presence of the variant or the treatment with the M channel blocker led to a large leftward shift of the relation between the number of AP and the amplitude of the current injected (#AP/I, Fig. 1Ac). The graphs were significantly different up to a current step of 220 pA. For current steps of larger amplitude the number of action potential in mutant cells was not different from that of wild type cells while it was significantly decreased in XE-991 treated wild type cells (Fig.1Abc). The initial and final frequencies of the discharge were also affected by the variant and the treatment with XE-991. A significant leftward shift of the initial frequency/ current relation (iF/I) was observed for the two groups of cells as well as the final frequency/current relation (fF/I) but for the latter the difference was significant for current steps up to160 pA. There was no statistical difference in the final frequency of the discharge between mutant and wild type cells for current steps of larger amplitude while the frequency was significantly lower for XE-991 treated wild type cells (Fig 1Ac). The decrease in the number and frequency of AP in mutant and XE treated wild type cells for current steps of strong amplitude were likely the consequence of the high input resistance of these 2 groups of cells leading to a much larger depolarization and to the progressive inactivation of voltage-gated Na^+^ channels.

**Figure 1:**
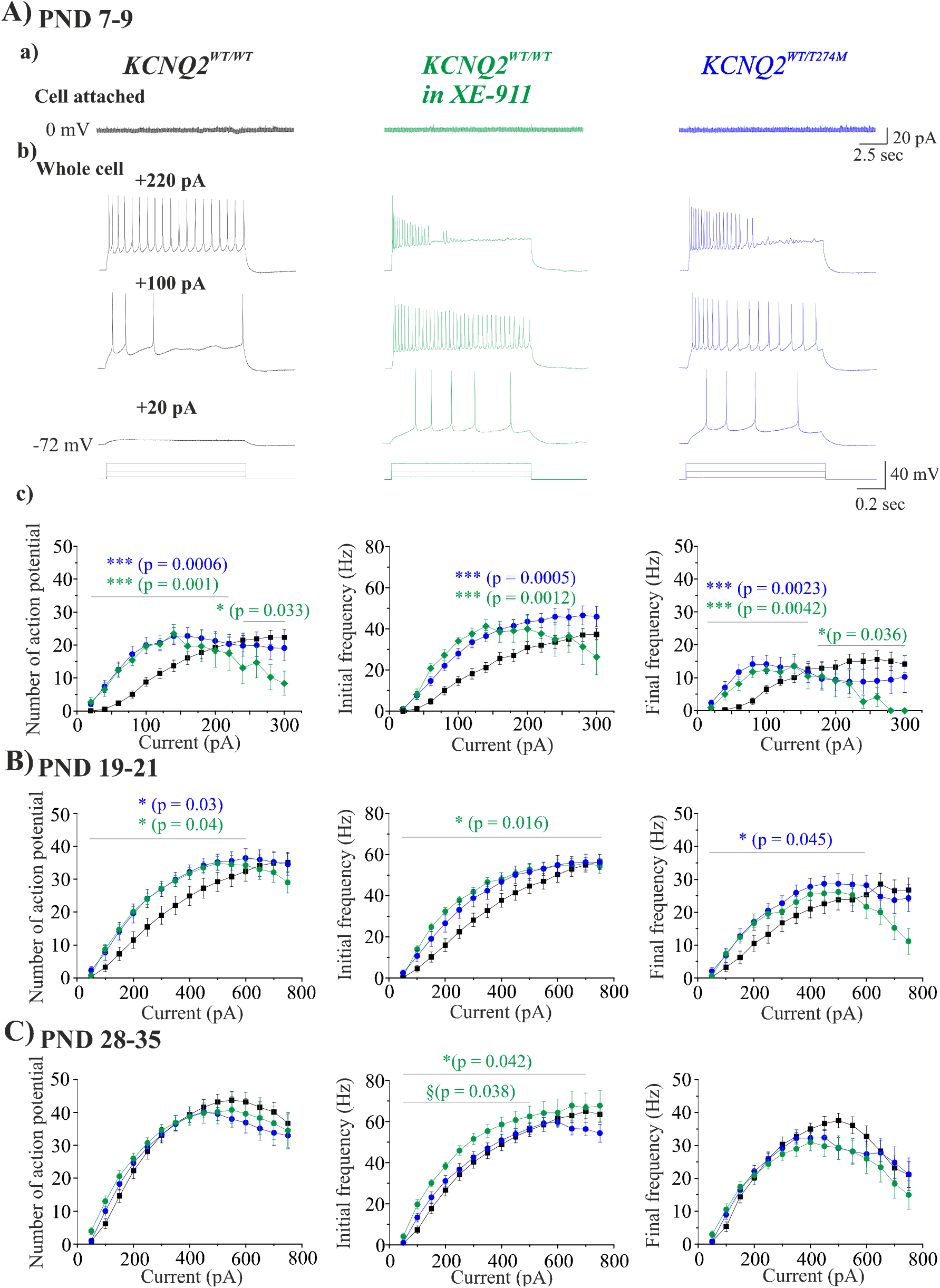
Effects of XE-991 and of the p.T274M variant on the discharge of developing layers II/III pyramidal cells and elicited by injection of current steps. **Aa)**Representative traces showing the recordings of pyramidal cells in cell-attached configuration in motor cortical slices from a *KCNQ2^WT/WT^* mouse (left traces, black), from a *KCNQ2^WT/WT^* mouse and superfused with 20 µM XE-991 (middle traces, green), from a *KCNQ2^WT/T274M^* mouse (left trace, blue) and aged one week (Post-Natal Days, PND 7-9). **Ab)** Same cells recorded in whole cell configuration and showing the voltage responses to the injection of three depolarizing current steps of 20, 100 and 220 pA applied during 1 sec. **Ac)** Graphs showing the quantification (mean ± S.E.M) at PND 7-9 of the number of action potentials elicited by the injection of depolarizing current steps (left graph), the initial frequency (measured during the first 200 msec of the steps, middle graph) and final frequency (measured during the last 200 msec of the steps, right graph) of the discharge. Black: wild-type cells (n = 17 cells, 2 animals); green: XE-991 treated wild type cells (n = 11 cells, 2 animals); blue: mutant cells (n = 17, 3 animals). **B)** Same quantifications as in Ac in cells from mice aged three weeks (PND 19-21). Black: wild-type cells (n = 20 cells, 4 animals); green: XE-991 treated wild type cells (n = 21 cells, 4 animals); blue: mutant cells (n = 22 cells, 4 animals). **C)** Same quantification of the three parameters in cells from mice aged four-five weeks (PND 28-35). Black: wild type cells (n = 27 cells, 5 animals); green: XE-991 treated wild type cells (n = 25 cells, 5 animals); blue: mutant cells (n = 25 cells, 4 animals). Statistics: * Comparison of *KCNQ2^WT/T274M^* (mutant cells) or *KCNQ2^WT/WT^* in XE-991(XE-991 treated wild type cells) versus *KCNQ2^WT/WT^* (wild type cells) ; § comparison of XE-991 treated wild type cells versus mutant cells. P values are indicated when differences were statistically significant (two-way Anova test)

**Table 1:** Intrinsic properties of pyramidal cells of the layer II/III in motor cortical slices from KCNQ2^WT/WT^ mice (black), in slices from KCNQ2^WT/WT^ mice superfused with XE-991 (20 µM, green), in slices from KCNQ2^WT/T274M^ mice (blue) aged one week, three weeks and 4-5 weeks. Statistics: * KCNQ2^WT/T274M^ or KCNQ2^WT/WT^ in XE-991 (XE-991 treated wild type cells) vs KCNQ2^WT/WT^ (mutant cells) vs KCNQ2^WT/WT^ (wild type cells) * : p< 0.05 ; ** : p< 0.01 ; *** p<0.001.

| **Layers II/III** | **PN7-9**<br>*KCNQ2*<sup>WT/WT</sup> ;<br>*KCNQ2*<sup>WT/WT</sup> in XE-991<br>*KCNQ2*<sup>WT/T274M</sup> | **PN19-21**<br>*KCNQ2*<sup>WT/WT</sup><br>*KCNQ2*<sup>WT/WT</sup> in XE-991<br>*KCNQ2*<sup>WT/T274M</sup> | **PN28-35**<br>*KCNQ2*<sup>WT/WT</sup><br>*KCNQ2*<sup>WT/WT</sup> in XE-991<br>*KCNQ2*<sup>WT/T274M</sup> |
| --- | --- | --- | --- |
| Cm (pF) | 67.6 ± 3.4<br>59.0 ± 4.1<br>59.3 ± 4.7 | 92.5 ± 4.6<br>112.7 ± 5.2<br>101.9 ± 5.9 | 110.3 ± 6.9<br>97.6 ± 6.0<br>106.0 ± 6.4 |
| V<sub>m</sub> (mV) | -65.9 ± 0.9<br>-67.8 ± 2.1<br>-66.3 ± 1.3 | -69.9 ± 0.9<br>-68.9 ± 1.2<br>-72.3 ± 1.1 | -73.7 ± 1.2<br>-70.2 ± 1.3<br>-73.4 ± 0.7 |
| R<sub>m</sub> (mOhm) | 324.2 ± 24.3<br>483.5 ± 33.1<sup>**</sup> (p = 0.0012)<br>560.0 ± 53.4<sup>**</sup> (p = 0.0017) | 169.2 ± 14.3<br>203.3 ± 12.3<br>202.7 ± 20.5 | 157.8 ± 15.4<br>229.3 ± 19.6<sup>**</sup> (p = 0.0033)<br>185.6 ± 18.1 |
| τ<sub>m</sub> (msec) | 25.6 ± 1.7<br>26.7 ± 3.2<br>26.5 ± 2.3 | 17.2 ± 1.1<br>21.1 ± 1.4<sup>*</sup> (p = 0.034)<br>20.7 ± 1.4 | 15.4 ± 0.7<br>20.7 ± 1.8<sup>*</sup> (p = 0.024)<br>16.7 ± 1.1 |
| **Action potential properties** |  |  |  |
| Rheobase (pA) | 323.5 ± 13.9<br>218.2 ± 8.7<sup>***</sup> (p < 0.0001)<br>216.9 ± 13.9<sup>***</sup> (p < 0.0001) | 420.0 ± 24.4<br>447.6 ± 24.3<br>428.6 ± 32.1 | 446.3 ± 34.4<br>361.4 ± 26.3<br>371.2 ± 31.0 |
| Threshold (mV) | -37.9 ± 0.7<br>-43.2 ± 0.8<sup>***</sup> (p < 0.0001)<br>-41.4 ± 0.6<sup>*</sup> (p = 0.017) | -38.1 ± 1.3<br>-42.9 ± 1.1<sup>§</sup> (p = 0.013)<br>-39.4 ± 1.2 | -46.4 ± 0.8<br>-47.8 ± 0.8<br>-46.9 ± 0.7 |
| Amplitude (mV) | 71.4 ± 1.4<br>76.1 ± 1.3<sup>*</sup> (p = 0.036)<br>72.1 ± 1.8 | 84.5 ± 2.1<br>90.5 ± 1.5<sup>*</sup> (p = 0.04)<br>86.8 ± 1.9 | 94.1 ± 1.1<br>95.4 ± 1.7<br>91.2 ± 1.8 |
| Overshoot (mV) | 35.3 ± 1.3<br>34.9 ± 1.4<br>33.5 ± 1.4 | 46.2 ± 1.2<br>47.8 ± 1.0<br>47.2 ± 1.3 | 49.5 ± 1.0<br>49.8 ± 1.5<br>46.3 ± 1.5 |
| Halfwidth (msec) | 1.1 ± 0.08<br>1.3 ± 0.12<br>0.9 ± 0.03<sup>*</sup> (p = 0.017) | 0.72 ± 0.03<br>0.82 ± 0.03<sup>*</sup> (p = 0.025)<br>0.83 ± 0.05 | 0.78 ± 0.03<br>0.91 ± 0.06<br>0.78 ± 0.03 |

We next examined the membrane response of the same cells to the injection of a slow ramp of current of 300 pA applied in 10 sec. This procedure should reveal much better the action of Kv7 channels which do not present voltage and time dependent inactivation contrary to many other ion channels. Several parameters of the response were analyzed. These included the resistance of the ramp measured between −72 mV and the action potential threshold (R_ramp_), the current required to evoke the first action potential (I_threshold_), the number of AP, the initial frequency during the first 200 msec of the discharge and the variation of voltage produced by the ramp (delta V). All these parameters were significantly affected in the same direction by the variant or following the inhibition of Kv7 channels (Fig. 2A). Compared to wild type cells (n = 17), the current threshold of both mutant cells (n = 17) and of XE-991 treated wild type cells (n = 11) was decreased, the AP membrane threshold more hyperpolarized (only for XE-991 treated cells), the initial frequency of the discharge was increased but the total number of AP was less due to a larger depolarization and the inactivation of voltage-gated Na^+^ channels.

**Figure 2:**
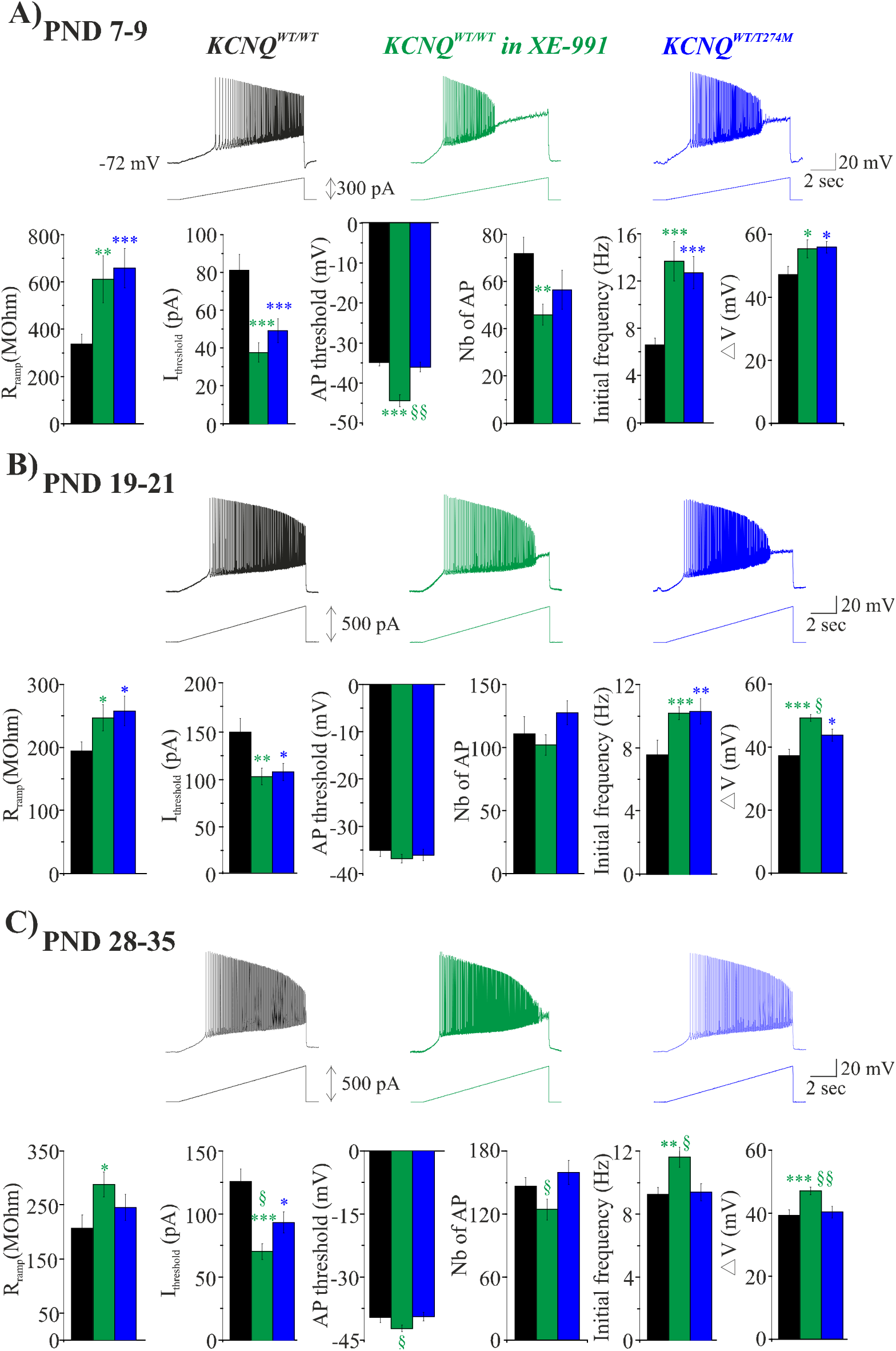
Effects of XE-991 and of the p.T274M variant on the voltage responses of developing layers II/III pyramidal cells to the injection of a ramp of current. **A)** Traces showing the response of pyramidal cells to the injection of a ramp of 300 pA in 10 sec recorded in motor cortical slices from *KCNQ2^WT/WT^* (black), *KCNQ2^WT/WT^* in presence of XE-991 (green) and *KCNQ2^WT/T274^* (blue) mice aged one week. Below traces are shown histograms representing the quantification (mean ± S.E.M) in the three group of cells of the resistance, the current and voltage threshold of first action potential generation, the number of action potential, the initial frequency (first 200 msec of the discharge) and the variation of the membrane potential produced by a ramp of current of 300 pA. Black: wild type cells (n = 17 cells, 2 animals); green: XE-991 treated wild type cells (n = 11 cells, 2 animals); blue: mutant cells (n = 17 cells, 3 animals). **B)** Same as in A in cells from mice aged three weeks. Analysis were performed on the voltage responses to the injection of a ramp of 500 pA in 10 sec. Black: wild-type cells (n = 20 cells, 4 animals); green: XE-991 treated wild type cells (n = 21 cells, 4 animals); blue: mutant cells (n = 22 cells, 4 animals). **C)** Same as in B in cells from mice aged 4-5 weeks. Black: wild type cells (n = 27 cells, 5 animals); green: XE-991 treated wild type cells (n = 25 cells, 5 animals); blue: mutant cells (n = 25 cells, 4 animals). Statistics: * comparison of *KCNQ2^WT/T274M^* (mutant cells) or *KCNQ2^WT/WT^* in XE-991 (XE-991 treated wild type cells) vs *KCNQ2^WT/WT^* (wild type cells) ; § comparison of XE-991 treated wild type cells vs mutant cells. *§ : p< 0.05 ; **,§§ : p< 0.01 ; ***,§§§ p<0.001 (unpaired Student t-test and Mann - Whitney test).

Therefore, at the end of the first postnatal week of life, the variant or the inhibition of M channels had the same consequences on the electrophysiological properties of layers II/III pyramidal cells. Both make cells more excitable in response to the injection of steps or a ramp of current. However neither the p.T274M variant nor XE-991 rendered cells spontaneously more excited.

At PND19-21, wild-type pyramidal cells (n = 20 cells/ 4 animals) had higher capacitance, twice lower input resistance and faster membrane time constant than at PND7-9 (table 1). Moreover, the current threshold to elicit an AP by the injection of steps or ramp of current was higher (Figs. 1B, 2B). Most of the effects of the variant and of XE-991 described at PND7-9 on membrane responses to current injection were maintained at three weeks. Like at PND7-9, in cell-attached configuration, the variant or the treatment with the blocker did not led pyramidal cells to spontaneously fire AP (n = 22 mutant cells/4 animals and 21 XE-991 treated wild type cells/4 animals, data not shown). In whole cell configuration, the responses of the cells to the injection of steps (50-750 pA) or to a ramp of current (500 pA in 10sec) were similarly and significantly increased compared to the responses of wild type cells (Figs. 1B,2B). The excitability of pyramidal cells in these 2 former groups of cells was significantly enhanced compared to wild type cells and observed for a wider range of current injected at PND7-9.

At PND28-35, the response of wild type pyramidal cells to current steps but not to the injection of slow ramp of current was stronger than at PND19-21 (Figs. 1C, 2C). #AP/I, iF/I and fF/I relations were steeper at PND 28-35 (n = 27 wild type cells/5 animals, Fig. 1C). The presence of the variant affected neither the discharge of the AP elicited by the injection of current steps nor the response of the cells to the injection of a slow ramp of current (n = 25 mutant cells/4 animals). In contrast, XE-991 (n = 25 cells/5 animals) produced a significant leftward shift of the iF/I relation, although the #AP/I relation was unchanged (Fig. 1C), and significantly affected in the same way than at PND19-21 most of the parameters of the membrane response to the injection of a slow ramp of current (Fig. 2C).

Therefore, the increase pyramidal cells excitability produced by the variant is limited to the three first postnatal week of life while the electrophysiological alterations produced by XE-991 is maintained from neonatal to juvenile stages at least.

### Layer V pyramidal cells are impacted at a later developmental stage by the variant than in layers II/III but also transiently

We next wondered if the variant and XE-991 impacted the excitability of pyramidal cells of other layers of the motor cortex and if these cells presented the same developmental sensitivity than described in layers II/III. We observed that pyramidal cells of the layer V presented some differences in the sensitivity to XE-991 and to the variant compared to pyramidal cells of layers II/III. In particular, at PND7-9, cells (n = 16 wild type cells/ 3 animals) were affected neither by the variant (n = 17 mutant cells/3 animals) nor by the Kv7 channel blocker (n = 11 XE-991 treated wild type cells/ 3 animals, Figs 3A, 4A and Table 2). At PND19-21, the variant increased significantly the firing of pyramidal cells elicited by injection of current steps and affected some of the parameters of the response elicited by the injection of a slow ramp of current (n = 18 wild type cells/5 animals and n = 20 mutant cells/ 5 animals) as it did in layers II/III but the effects of the variant were of lower magnitude than those produced by XE-991 ( n = 19 XE-991treated wild type cells/ 4 animals, Figs. 3B, 4B). At PND28-35, like for pyramidal cells of layers II/III, the variant had no more consequences on layer V pyramidal cells whereas XE-991 increased neuronal discharge evoked by current steps and affected significantly most of the parameters of the responses elicited by the injection of a ramp of current (n = 22 wild type cells/ 5 animals, n = 18 XE-991-treated wild type cells/ 5 animals, n = 20 mutant cells/ 4 animals, Figs. 3C, 4C).

**Figure 3.**
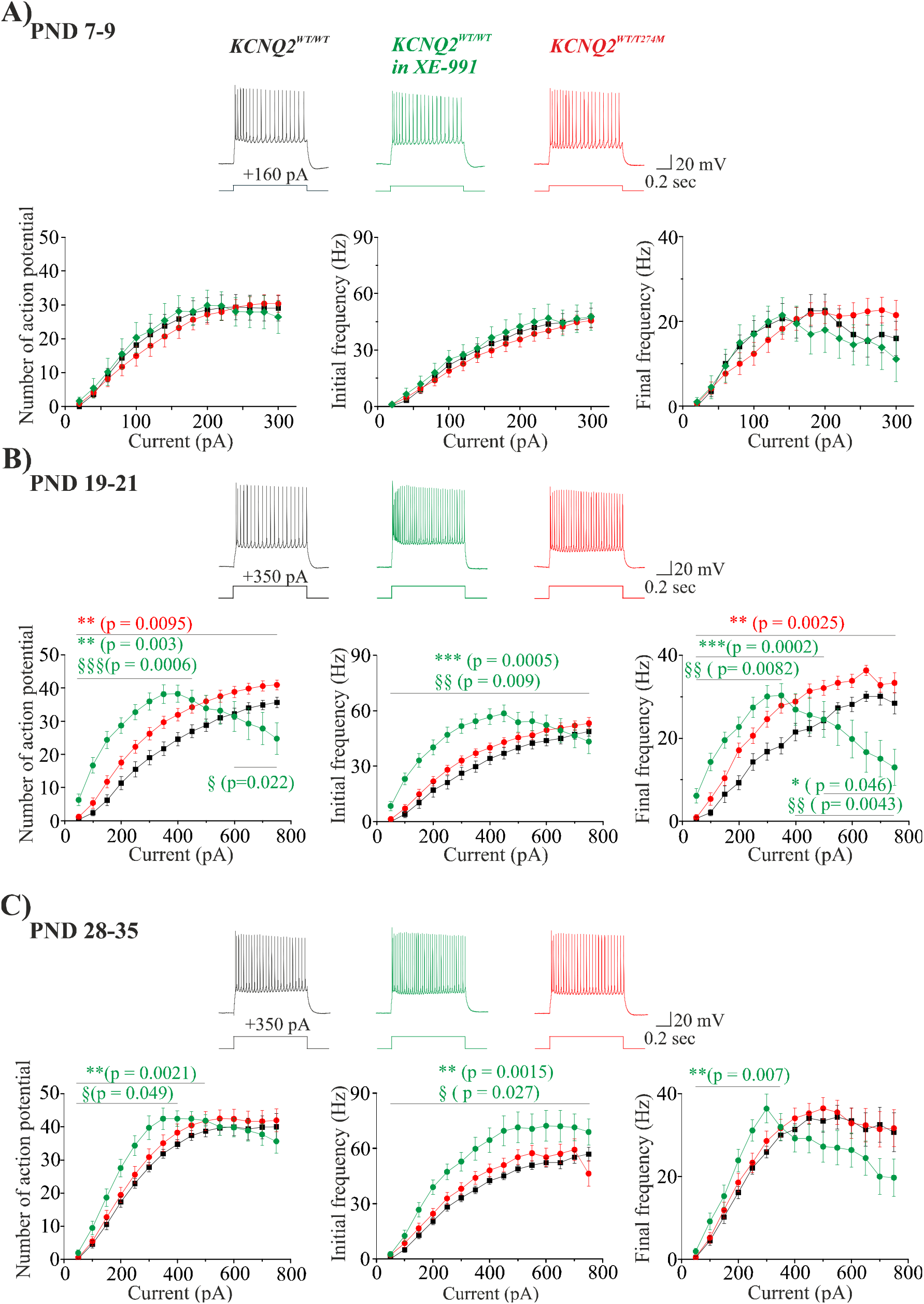
Effects of XE-991 and of the p.T274M variant on the discharge of developing pyramidal cells of the layer V elicited by the injection of current steps **A)** Representative traces showing the voltage responses to the injection of current step of 160 pA for 1 sec in layer V pyramidal cells in motor cortical slices from *KCNQ2^WT/WT^* (black), *KCNQ2^WT/WT^* in presence of XE-991 (green), *KCNQ2^WT/T274M^* (red) mice aged one week (PND 7-9). Below traces are shown the quantification (mean ± S.E.M) of the number of action potentials (left graph) elicited by the injection of depolarizing current steps, the initial and final frequencies of the discharge (middle and right graphs respectively) in pyramidal cells from animals aged one week (PND 7-9). Black: wild-type cells (n = 16 cells, 3 animals); green: XE-991 treated wild type cells (n = 11 cells, 3 animals); red: mutant cells (n = 17, 3 animals). **B)** Same as in A but for animals aged 3 weeks (PND 19-21). Traces depicted show voltage responses of pyramidal cells to the injection of current step of 350 pA. Quantifications have been done on 18 wild-type cells, 5 animals (black), 19 XE-treated wild type cells, 4 animals (green); 20 mutant cells, 5 animals (red). **C)** Same as in B but from animals aged 4-5 weeks (PND28-35). Quantifications have been done on 22 wild type cells, 5 animals (Black), 18 XE-991 treated wild type cells, 5 animals (green), 20 mutant cells, 4 animals (red) Statistics: *comparison of XE-991 treated wild type cells or mutant cells vs wild type cells ; § comparison of XE-911 treated wild type cells vs mutant cells. P values are indicated when differences were statistically significant (two-way Anova test).

**Figure 4:**
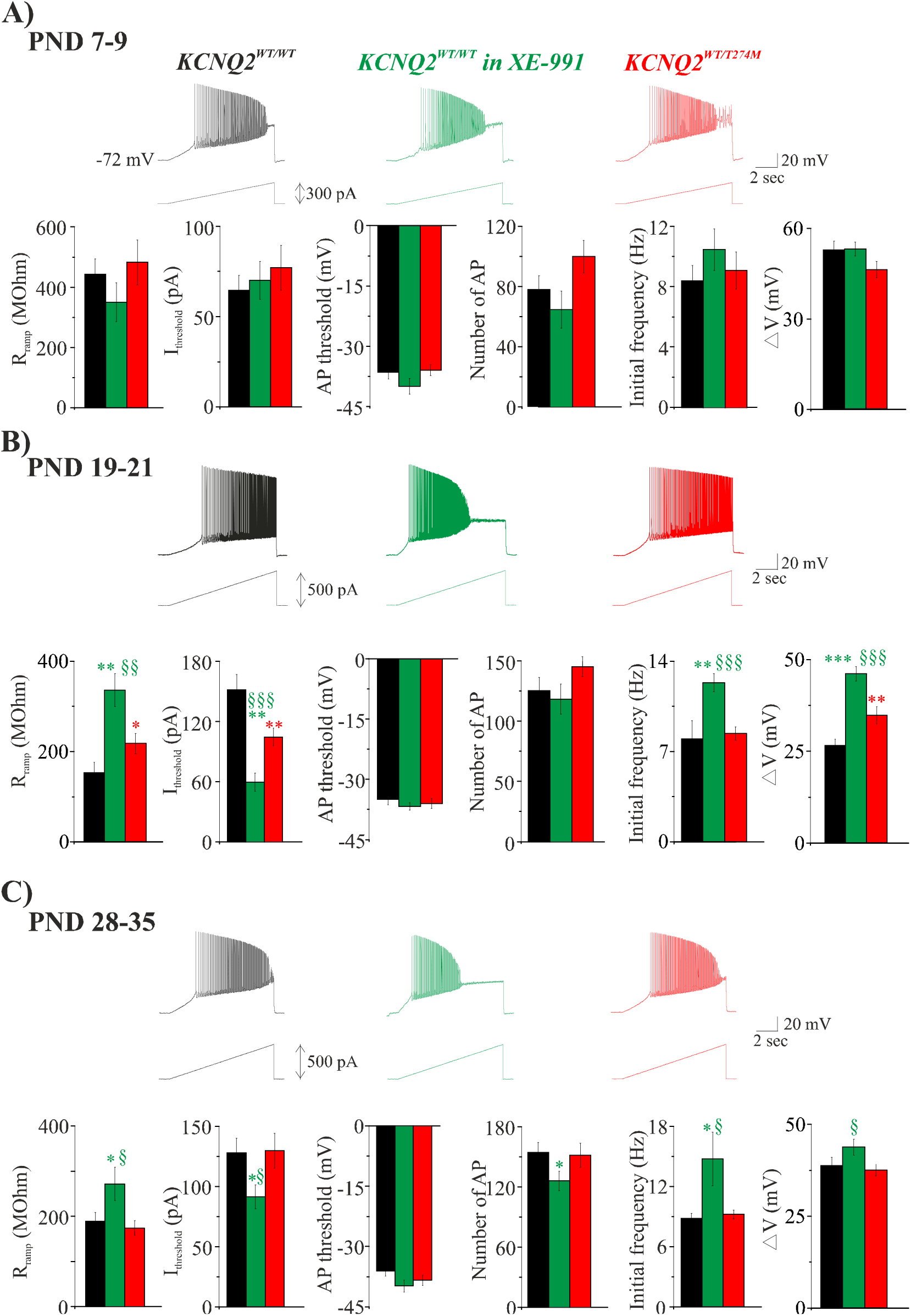
Effects of XE-991 and of the p.T274M variant on the voltage responses of developing pyramidal cells of the layer V to the injection of a ramp of current. **A)** Traces showing the response of pyramidal cells to the injection of a ramp of 300 pA in 10 sec recorded in motor cortical slices from *KCNQ2^WT/WT^* (black), *KCNQ2^WT/WT^* in presence of XE-991 (green) and *KCNQ2^WT/T274^* (red) mice aged one week. Below traces: Histograms representing the quantification (mean ± S.E.M) in the three group of cells of the resistance, the current and voltage threshold of first action potential generation, the number of action potential, the initial frequency (first 1sec of the discharge) and the variation of the membrane potential produced by the ramp of current of 300 pA. Black: wild type cells (n = 16 cells, 3 animals); green: XE-991 treated wild type cells ( n = 11 cells, 3 animals); red: mutant cells (n = 17 cells, 3 animals). **B)** Same as in A in cells from mice aged three weeks. Analysis were performed on the voltage response elicited by the injection of a ramp of 500 pA in 10 sec. Black: wild-type cells (n = 18 cells, 5 animals); green: XE-991 treated wild type cells (n = 19 cells, 4 animals); red: mutant cells (n = 20 cells, 5 animals). **C)** Same as in B in cells from mice aged 4-5 weeks. Black: wild type cells (n = 22 cells, 5 animals); green: XE-991 treated wild type cells (n = 18 cells, 5 animals); red: mutant cells (n = 20 cells, 4 animals).

**Table 2:** Intrinsic properties of pyramidal cells of the layer V in motor cortical slices from KCNQ2^WT/WT^ mice (black), in slices from KCNQ2^WT/WT^ mice superfused with XE-991 (20 µM, green), and in slices from KCNQ2^WT/T274M^ mice (red) aged one week, three weeks and 4-5 weeks.

| **Layer V** | **P7-P9**<br>*KCNQ2<sup>WT/WT</sup>*<br>*KCNQ2<sup>WT/WT</sup> in XE-991*<br>*KCNQ2<sup>WT/T274M</sup>* | **P19-P21**<br>*KCNQ2<sup>WT/WT</sup>*<br>*KCNQ2<sup>WT/WT</sup> in XE-991*<br>*KCNQ2<sup>WT/T274M</sup>* | **P28-P35**<br>*KCNQ2<sup>WT/WT</sup>*<br>*KCNQ2<sup>WT/WT</sup> in XE-991*<br>*KCNQ2<sup>WT/T274M</sup>* |
| --- | --- | --- | --- |
| Cm (pF) | 52.9 ± 6.4<br>57.0 ± 4.4<br>62.3 ± 8.7 | 108.8 ± 5.5<br>99.9 ± 11.0<br>119.6 ± 5.5 | 111.8 ± 6.1<br>101.1 ± 10.1<br>114.1 ± 6.0 |
| Vm (mV) | -63.1 ± 1.4<br>-63.3 ± 1.3<br>-61.2 ± 1.9 | -69.4 ± 1.1<br>-68.1 ± 1.6<br>-71.1 ± 0.9 | -68.9 ± 1.5<br>-68.9 ± 1.4<br>-69.2 ± 1.0 |
| Rm (mOhm) | 515.6 ± 60.1<br>451.0 ± 65.5<br>427.3 ± 63.8 | 126.6 ± 20.3<br>262.7 ± 31.6<sup>**</sup> (p = 0.0013)<br>145.5 ± 14.7 | 158.5 ± 13.3<br>208.9 ± 30.1<br>160.6 ± 18.5 |
| τ<sub>m</sub> (msec) | 22.7 ± 2.1<br>23.7 ± 2.3<br>25.7 ± 2.4 | 16.5 ± 2.3<br>21.0 ± 2.0<br>17.0 ± 1.5 | 16.6 ± 1.6<br>17.7 ± 2.5<br>15.1 ± 1.3 |
| **Action potential Properties** |  |  |  |
| Rheobase (pA) | 233.3 ± 21.4<br>254 ± 21.1<br>233.3 ± 22.3 | 457.9 ± 31.6<br>357.1 ± 24.7<sup>*</sup> (p = 0.017)<br>507.9 ± 31.1 | 484.1 ± 28.9<br>412.4 ± 30.5<br>464.3 ± 36.9 |
| Threshold (mV) | -37.9 ± 1.1<br>-39.2 ± 2.1<br>-37.9 ± 1.1 | -47.9 ± 1.3<br>-46.3 ± 1.0<br>-45.6 ± 1.2 | -43.0 ± 1.2<br>-44.0 ± 1.2<br>-44.9 ± 0.9 |
| Amplitude (mV) | 75.9 ± 2.5<br>73.3 ± 2.3<br>80.6 ± 1.5 | 90.8 ± 1.3<br>89.0 ± 1.7<br>90.2 ± 1.6 | 90.5 ± 1.4<br>89.6 ± 1.4<br>89.3 ± 1.6 |
| Overshoot (mV) | 37.9 ± 1.2<br>36.2 ± 2.8<br>43.1 ± 2.5 | 45.1 ± 1.1<br>43.6 ± 1.4<br>46.3 ± 2.0 | 49.4 ± 1.4<br>46.3 ± 1.1<br>45.8 ± 1.2 |
| Halfwidth (msec) | 1.2 ± 0.1<br>1.5 ± 0.1<br>1.0 ± 0.07 | 0.70 ± 0.05<br>0.83 ± 0.03<sup>**</sup> (p = 0.0013)<br>0.75 ± 0.03 | 0.77 ± 0.04<br>0.70 ± 0.02<br>0.78 ± 0.04 |

Together these data showed that the action of the p.T274M variant on neuronal excitability in layers II/III and V is developmentally regulated and that the relative contribution of Kv7/M channels on pyramidal cells discharge in layers II/III and V is not equivalent.

### Developmental effect of the p.T274M variant on M current in pyramidal cells of layers II/III and V

The increase in neuronal excitability produced by the variant is likely the consequence of a reduction in M current. Previous studies performed in heterologous cells showed that the p.T274M variant reduced by ∼60 % M current carried by heteromeric Kv7.2/Kv7.3 channels (Orhan et al., 2014) and by ∼50% in interneurons of the locomotor CPG region of the spinal cord (Verneuil et al., 2021). We therefore analyzed the impact of the variant on M current in pyramidal cells of the layers II/III and V at the same three developmental stages from PND 7-9 to PND 28-35.

M current was analyzed using the standard Im deactivation protocol described by Adams et al. (1982). Hyperpolarizing voltage steps down to −90 mV in 10 mV increments (1 sec duration) from a membrane potential of +10 mV were applied (see Material and Methods). These hyperpolarizing steps led to an inward current relaxation which amplitude calculated by subtracting steady state current (Iss) from instantaneous (Iins) current levels at each membrane potentials increased from 0 to −30 mV and then decreased at more hyperpolarized membrane potentials. The intersection of the Iins and Iss currents yielded apparent reversal potentials ranging from ∼-62 mV to −75 mV in wild type cells of the layers II/III and V depending of the developmental stage (see below). First, in order to confirm that the current relaxation corresponded to the closing of Kv7 channels and the deactivation of M current, we recorded six wild type pyramidal cells in layers II/III of motor cortical slices of the left hemisphere in absence of XE-991 and six wild type pyramidal cells in layers II/III of motor cortical slices of the right hemisphere from same animal aged 3 weeks and in presence of XE-991 (20 µM). The presence of the Kv7 channels blocker led to a lower holding current at +10 mV and to a strong reduction (by ∼85%) of the current relaxation at all membrane potentials (Fig. 5). These data demonstrated that the inward relaxation was mediated by Im.

**Figure 5:**
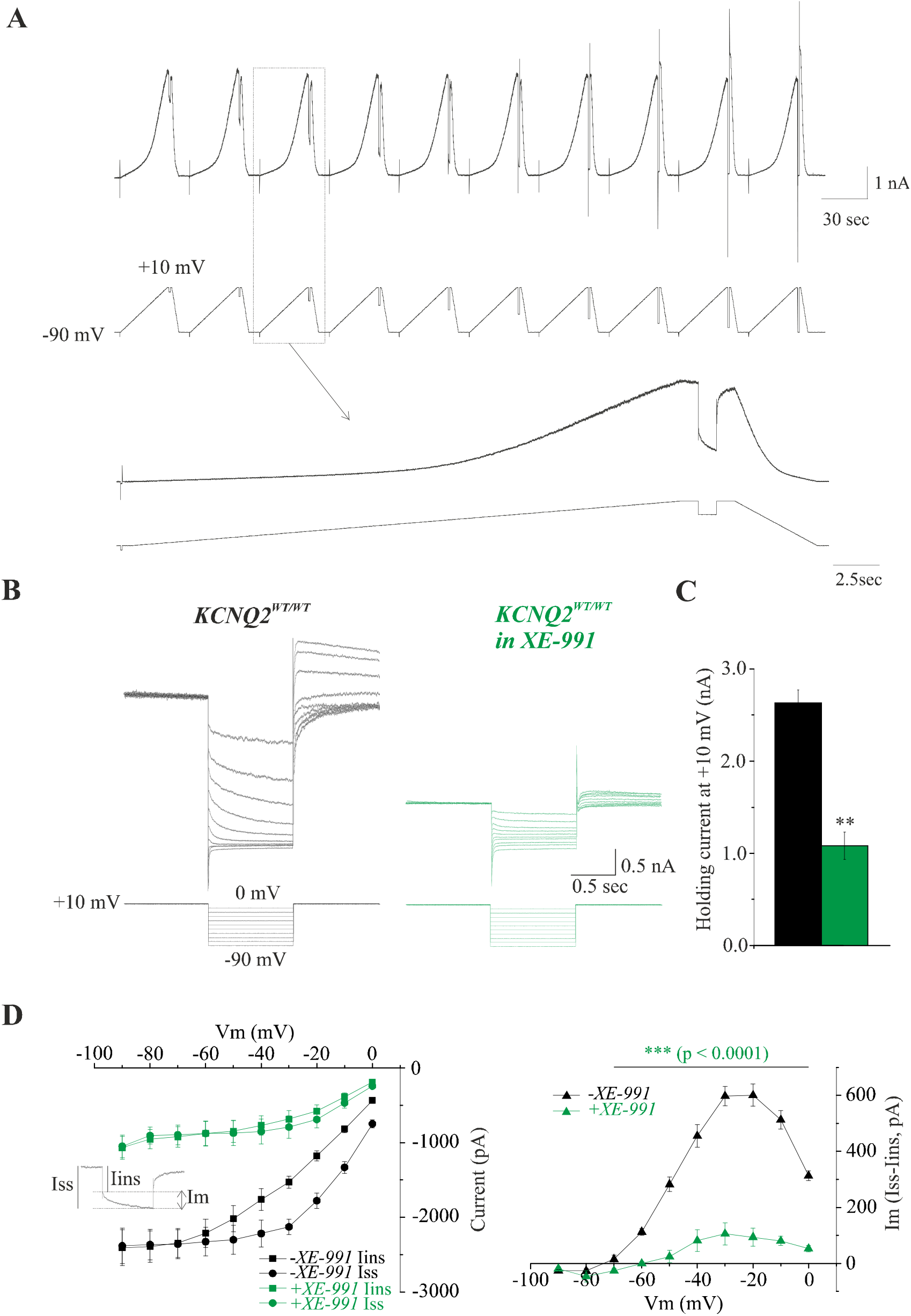
Characterization of M current in pyramidal cells. **A)** Experimental protocol used to isolate M current. Slow ramps of voltage are applied from −90 mV to reach +10 mV in 30 sec to activate Kv7/M channels. Cells were maintained for 1 sec at this membrane potential and hyperpolarizing voltage steps down to −90 mV (in 10 mV increments) are applied to deactivate the channels leading to an inward current relaxation. Bottom traces show at a higher time scale an example of current response to a slow ramp of voltage followed by an hyperpolarizing voltage step to −20 mV. **B)** Isolated current responses to hyperpolarizing voltage steps command applied from +10 mV down to −90 mV recorded in a wild type layers II/III pyramidal cells in left motor cortical slice at PND 20 in absence of XE-991 and from another pyramidal cell in right motor cortical slice from same preparation in presence of XE-991 (20 µM). **C)** Histograms showing the mean holding current measured at +10 mV in cells recorded in absence ( n = 6 cells, black) and in presence of XE-991 (n = 6 cells, green). **D)** Left graph: Instantaneous (Iins, square symbols) and steady-state (Iss, circle symbols) current (I) / voltage (V) relations of wild type cells in absence (black symbols) and in presence of XE-991 (green symbols). The apparent reversal potential of the current is the intersection between the two curves. Right graph: Graph showing the amplitude of the inward current relaxation measured as difference between instantaneous (Iin) and steady-state (Iss) levels at the onset and at the end of each hyperpolarizing voltage step command. Current relaxation represents the deactivation of Kv7/M channels.

M current was present in pyramidal cells of layers II/III at PND7-9 and was sensitive to the p.T274M variant. We measured current density (I) and calculated chord conductance (G) at each membrane potentials (I/V and G/V relations respectively). Compared to wild-type cells, both I/V and G/V relations of mutant cells were significantly smaller (n = 16 wild type cells/3 animals and n = 17 mutant cells/4 animals, Fig 6A). The apparent reversal potential (intersection of Iss and Iins) was, on average not significantly, affected by the variant (-66.3 ± 2.2 mV and −61.9 ± 2.9 mV in wild type and mutant cells respectively, p = 0.21 unpaired t-test). At PND19-21, current density and chord conductance measured in wild type cells were respectively ∼4 and ∼5 times higher than at PND 7-9 (n = 17 wild type cells/4 animals). The apparent reversal potential was also significantly more hyperpolarized (-75.6 ± 1.3 mV, p = 0.0007 unpaired t-test). Both I/V and G/V relations were significantly smaller in mutant cells compared to wild type cells at PND19-21 without any consequences on reversal potential ( −72.6 ± 1.6 mV, p =0.16 unpaired t-test, n = 16 mutant cells/4 animals, Fig.6B). At PND 28-35, I/V and G/V relations in wild type cells were significantly smaller than in wild type cells at PND19-21 (p = 0.017 and p = 0.015 respectively, two-way Anova test between 0 and −60 mV, n = 16 wild type cells/3 animals, Fig. 6B,C), while both relations were similar in mutant cells at these two developmental stages (p = 0.39 and p = 0.41 respectively, two-way Anova test; n = 16 mutant cells/4 animals at PND 28-35, Fig. 6B,C). Reversal potentials in wild and mutant cells were −74.3 ± 1.5 mV and −74.4 ± 2.2 mV respectively. Importantly at PND28-35, I/V and G/V relations in both group of cells were not different (p = 0.42 and p = 0.3 respectively Two-way Anova test, Fig 6C). Finally the p.T274M variant did not alter the deactivation time constant measured between 0 mV and −60 mV and at the three developmental stages. There were also not significant variations of the time constant values measured at each membrane potential from PND7-9 to PND 28-35.

**Figure 6:**
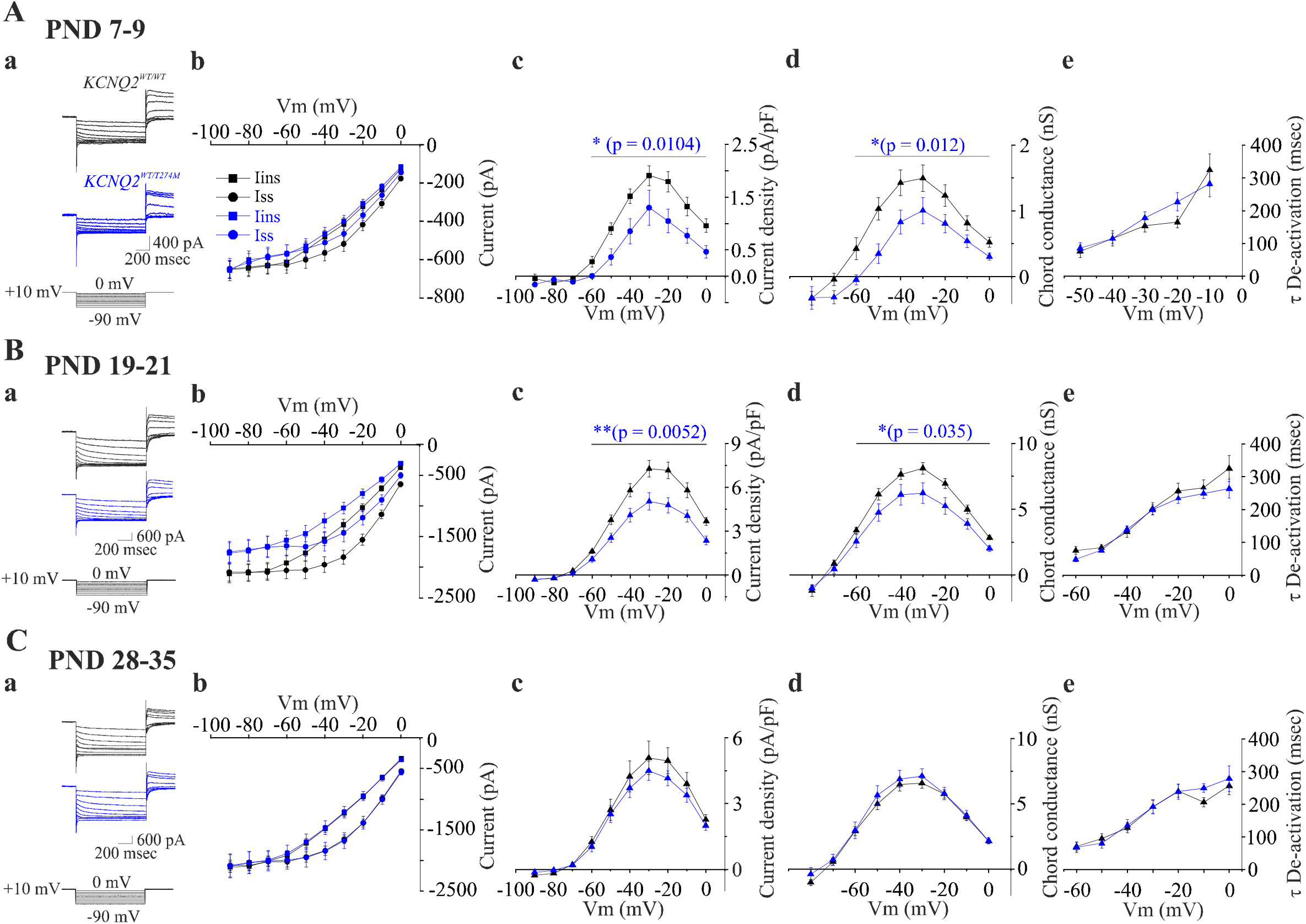
M current analysis in layers II/III pyramidal cells from developing *KCNQ2^WT/WT^* and *KCNQ^WT/T274M^* mice **Aa)** Representative current responses to hyperpolarizing voltage steps in a wild type pyramidal cell (black) and a mutant pyramidal cell (blue) recorded at PND 8 and PND 9 respectively. **Ab)** Instantaneous (Iins) and steady-state current (Iss) – voltage relations measured in wild-type cells (n = 16, 4 animals, black symbols) and in mutant cells (n = 16 cells, 4 animals, blue symbols). **Ac-e)** Plots of M current density **(c)**, chord conductance **(d)** and deactivation time constant (e) versus membrane potentials of wild type and mutant pyramidal cells from animals aged one week. **B)** Same as in A for wild type and mutant pyramidal cells from animals aged 3 weeks ( n= 17 wild type cells, 4 animals and n = 16 mutant cells, 4 animals). Traces in Ba are from wild type and mutant cells recorded at PND 20 and PND 21 respectively. **C)** Same measurements performed at PND28-35 in 16 wild-type cells (3 animals) and 16 mutant cells (4 animals). Traces in **Ca** are from a wild type and mutant cells recorded at PND 29 and PND 30 respectively. P values are indicated when comparison of the plots of wild-type with mutant cells were statistically significant (two-way Anova test performed between 0 and −60 mV).

In wild type pyramidal cells of layer V, current density and chord conductance increased from PND 7-9 (n = 16 wild type cells/4 animals) to PND 19-21 (n = 16 wild type cells/4 animals) by ∼4 and ∼9 fold respectively with no further increase at PND 28-35 (n = 17 wild type cells/ 5 animals, Fig. 7A,B). However current density and chord conductance at PND7-9 and PND19-21 were significantly smaller than in layers II/III at the same two developmental stages (at PND 7-9, p = 0.0002 and p = 0.0004 for I/V and G/V respectively, two-way Anova test; at PND 19-21, p < 0.0001 and p = 0.0033 for I/V and G/V respectively, two-way Anova test). At PND 28-35, I/V and G/V relations were similar in layer V and layers II/III (p = 0.11 and p = 0.28 for I and G respectively two-way Anova test). As shown in figure 7, in layer V, M current density and chord conductance were significantly reduced by the p.T274M variant only at PND19-21 (n = 16 mutant cells, 3 animals).

**Figure 7:**
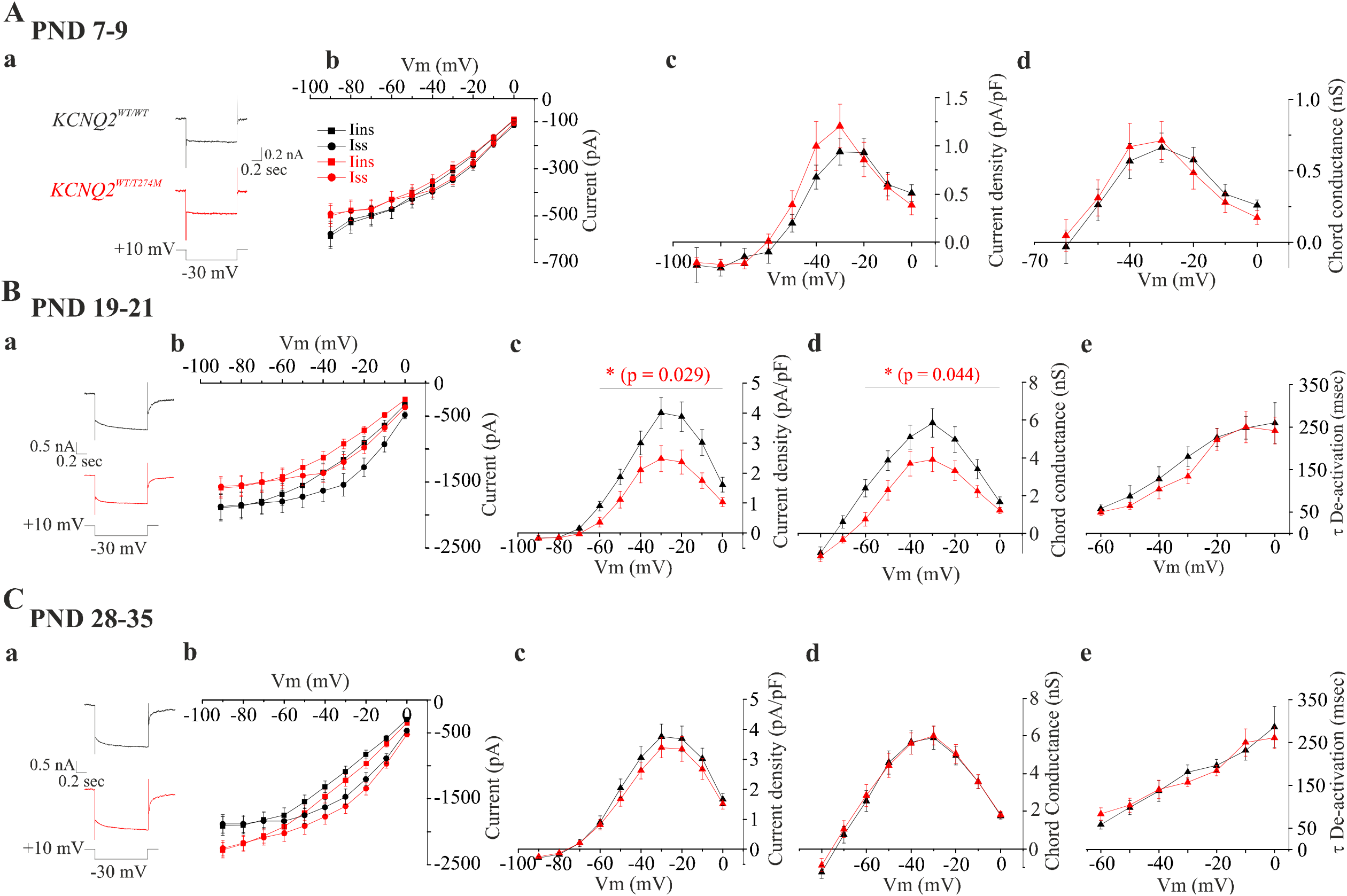
M current analysis in pyramidal cells of the layer V from developing *KCNQ2^WT/WT^* and *KCNQ^WT/T274M^* mice **Aa)** Representative current traces to an hyperpolarizing voltage step to −30 mV in a wild type pyramidal cell (black) and a mutant pyramidal cell (red) recorded at PND 7 and PND 9 respectively. **Ab)** Instantaneous (Iins) and steady-state current (Iss) – voltage relations measured in wild-type cells (n = 16, 4 animals, black symbols) and in mutant cells (n = 16 cells, 4 animals, red symbols). **Acd)** Plots of M current density **(c)** and chord conductance **(d)** versus membrane potentials of wild type and mutant pyramidal cells from animals aged one week. Measurement of deactivation time constant was not done because of the very small amplitude of the currents in both wild type and mutant pyramidal cells at this developmental stage. **B)** Same as in **A** for wild type and mutant pyramidal cells from animals aged 3 weeks ( n= 16 wild type cells, 4 animals and n = 16 mutant cells, 3 animals). Traces in **Ba** are from wild type and mutant cells recorded at PND 20 and PND 21 respectively. **Be)** Plot of deactivation time constant vs membrane potentials for wild-type and mutant cells. **C)** Same measurements performed at PND28-35 in 17 wild-type cells (5 animals) and 17 mutant cells (4 animals). Traces in **Ca** are from a wild type and mutant cells recorded at PND 29 and PND 30 respectively.

Together, these data showed that the p.T274 variant exerted a loss of function effect on Kv7/M channels in neocortical pyramidal cells and as for the analysis of neuronal excitability, the effect is developmentally regulated and no more observed in animals aged 4-5 weeks. Moreover these data showed that the expression of M current is stronger in layers II/III than in layer V until post-weaning stage but reached a similar level at a juvenile stage.

### Spontaneous network-driven events are increased in motor cortical slices from *KCNQ2^WT/T274M^* mice

We next analyzed spontaneous synaptic activity mediated by GABA and glutamate receptors at the same 3 developmental periods in layers II/III and V in motor cortical slices from *KCNQ2^WT/WT^* and *KCNQ2^WT/T274M^* mice. For this purpose pyramidal cells located in these layers were recorded in cell-attached and in whole-cell configurations in voltage clamp mode using patch pipettes filled with a Cs-gluconate solution (see material and methods). Postsynaptic currents mediated by GABA_A_ receptors (GABAR-PSCs) were recorded at 0 mV; the reversal potential of glutamate, while those mediated by glutamate receptors (GluR-PSCs) were recorded at the reversal potential of GABA estimated ∼-70 mV in our recording conditions.

In cell-attached configuration, we did not detected any spontaneous action potentials in wild-type and mutant cells of layers II/III and V recorded between PND7 to PND 35 (Fig.8A). Cells were still silent at their resting membrane potential although a “rippling” of the current in cell-attached configuration could be observed (Fig. 8B). In whole-cell configuration spontaneous postsynaptic events recorded in both populations of cells and at the three developmental periods were mainly mediated by GABA_A_ receptors. The frequency of these events was 3-10 times higher than GluR-PSCs (Fig.9). In addition, the activity recorded at 0 mV included recurrent outward currents with a magnitude of few hundreds of pA and lasting between 0.5-2 sec (Figs.8A,B and 9A). These large events were network activities resulting from the periodic summation of GABAergic postsynaptic currents. Indeed: i) their frequency did not change with the membrane potential; ii) they reversed polarity at the reversal potential of GABA_A_ receptors; iii) they were insensitive to NBQX/APV (n = 4/4 cells); iv) they were fully blocked by gabazine (n = 4/4 cells) (Fig.8B). These recurrent activities (Recurrent GABAergic Network Activity, RGNA) were more often observed in wild-type pyramidal cells of layers II/III than of the layer V and recorded until PND19-21. They were absent in cells from mice aged 4-5 weeks (Fig. 9 C,F).

**Figure 8:**
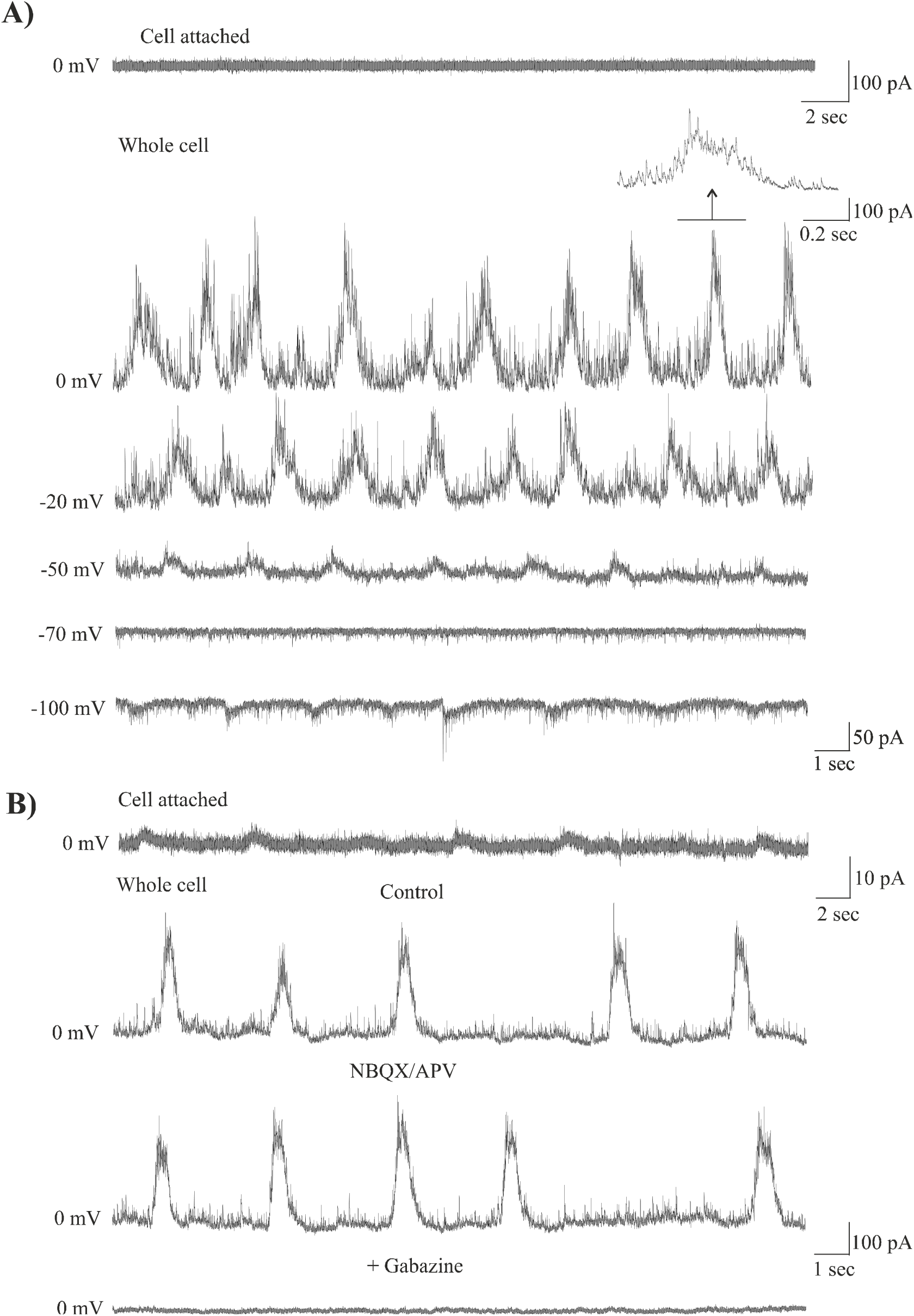
Recurrent GABAergic network activity in developing motor cortical slices**A)** Upper trace: pyramidal cell of layers II/III from a *KCNQ2^WT/T274M^* mouse aged 3 weeks and recorded in cell-attached configuration. Lower traces: same cell recorded in whole cell configuration. The spontaneous activity is characterized by the presence of recurrent large outward current reversing polarity at −70 mV and which frequency is insensitive to the membrane potential. **B)** Other pyramidal cell of the layers II/III from a *KCNQ2^WT/T274M^* mice aged 3 weeks and recorded in cell-attached (note the presence of small oscillation of the current) and in whole cell configurations. The traces show RGNA recorded in control, in the presence of NBQX (10 µM) and D-APV (40 µM), and in the presence of gabazine (5 µM).

**Figure 9:**
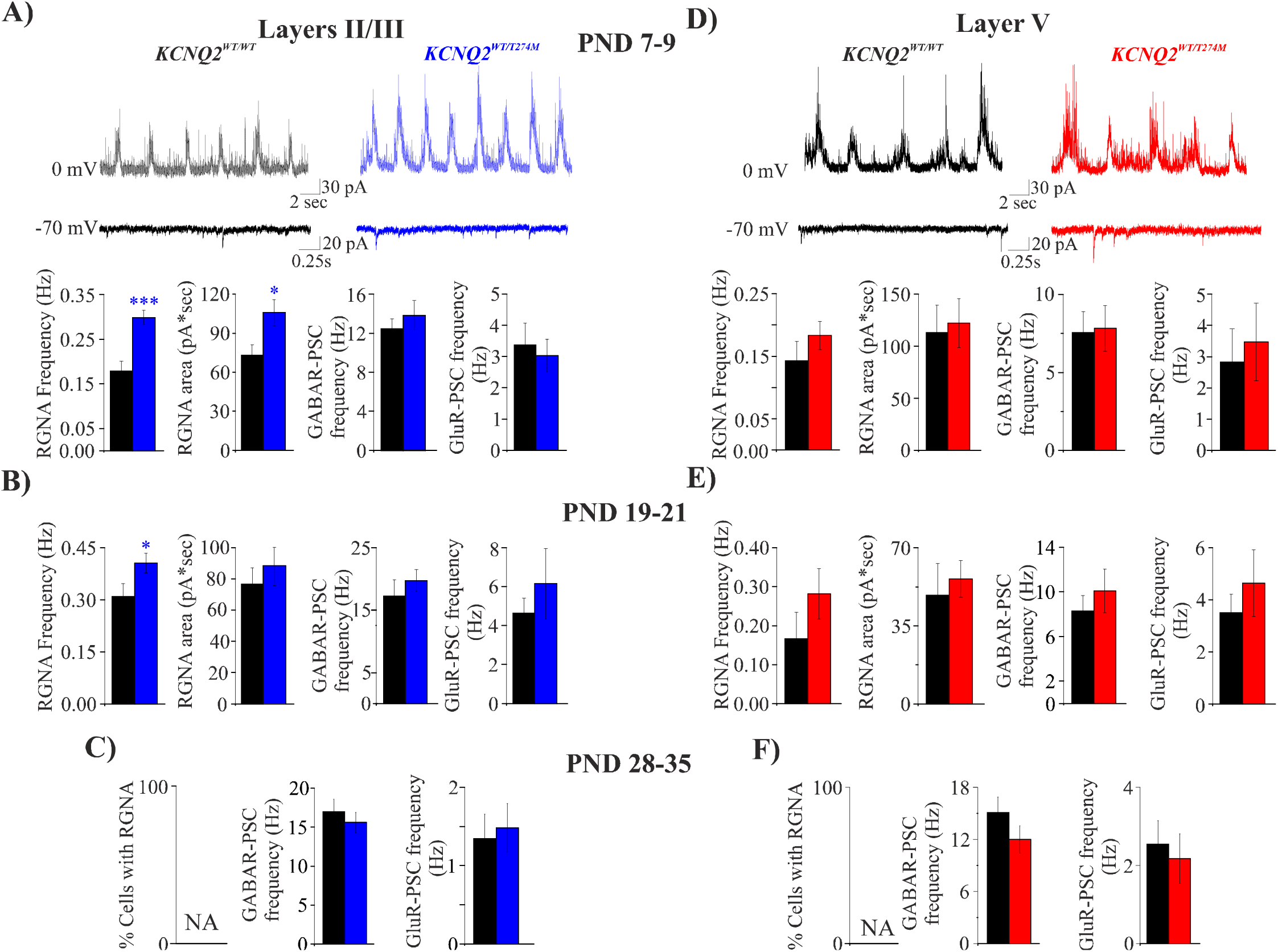
Effect of the p.T274M variant on spontaneous activities recorded in developing pyramidal cells of the layers II/III and V **A,D)** Representative traces at PND 7-9 showing spontaneous activities recorded at 0 mV and −70 mV in wild type and mutant pyramidal cells of the layers II/III (**A**, black and blue traces respectively) and of the layer V (**D**, black and red traces). GABAR-PSCs are recorded at 0 mV, GluR-PSCs are recorded at −70 mV. Below traces are shown the quantification (mean ± S.E.M) of the frequency of RGNA, the area of the outward current underlying RGNA, the total frequency of spontaneous GABAR-mediated post-synaptic currents (events in and out RGNA) and the frequency of GluR-mediated post-synaptic currents recorded in pyramidal cells at PND7-9. **B,E)** Same quantifications in layers II/III and V pyramidal cells from mice aged three weeks (PND19-21). **C,F)** Same quantifications in cells from mice aged 4-5 weeks (PND 28-35). RGNA were not observed (NA, not available). Statistics: * p< 0.05 ; ** : p< 0.01 ; ***p<0.001 (unpaired Student t-test).

At PND7-9, RGNA were recorded in 17/17 cells of layers II/III (3 animals), occurring at frequency ranging between 0.05 to 0.33 Hz and in 11/14 of cells of the layer V (4 animals) at frequency ranging between 0.03 to 0.33 Hz (Fig. 9A,D). At PND19-21, RGNA were recorded in 15/19 cells of layers II/III (3 animals) and at a higher frequency (ranging between 0.1 to 0.6 Hz) but only in 4/14 cells of the layer V (3 animals) at frequency ranging between 0.02 to 0.3 Hz (Fig. 9B,E). RGNA were not observed in upper and deep layers in animals aged 4-5 weeks (layers II/III: n = 28 cells, 4 animals; layer V: n= 22 cells, 3 animals, Fig. 9C,F).

The p.T274M variant had consequences on RGNA in layers II/III. In these layers, at PND 7-9, the frequency of RGNA as well as the area of outward currents underlying RGNA in mutant pyramidal cells were significantly increased compared to RGNA recorded in wild type pyramidal cells (n = 19 mutant cells, 4 animals, Fig. 9A). A small but significant increase in RGNA frequency but not of the area was also observed in mutant pyramidal cells at PND19-21 (n = 19 cells, 4 animals, Fig.9B). In addition RGNA were recorded in 100 % of the mutant cells but in 79% of wild type cells. However the frequency of GABAR-PSC (events in and out RGNA) was not significantly different suggesting that in the knock-in mice most of GABAR-PSC were concentrated in the RGNA. The frequency of GABAR-PSC was also unaffected at PND28-35 (n = 28 mutant cells, 4 animals, Fig. 9C). There was also no significant difference in the frequency of GluR-PSC between PND7 to PND 35 (Fig.9).

Ongoing synaptic activity recorded in the layer V showed some differences with that recorded in layers II/III. The occurrence of RGNA in wild type cells of the layer V and the number of cells in which they have been recorded were less than in layers II/ III. This was particularly obvious at PND19-21 where in the layer V, RGNA were recorded in only 30% of cells (4 out of 14 cells). Moreover the frequency of RGNA in layer V was on average twice lower than in layers II/III (Fig.9E). RGNA in the layer V appeared also to be less sensitive to the variant. Thus, at both PND7-9 and PND 19-21, there was no significant differences in the frequency of RGNA between wild-type cells (n = 14 wild type cells/ 3 animals and 14 wild type cells/3 animals, respectively) and mutant cells (n = 14 mutant cells/ 3 animals and n = 18 mutant cells/ 4 animals respectively, Fig.9D,E). However, at PND19-21, RGNA was recorded in more mutant cells (10 out of 18 cells) than in wild type cells (4 out of 14 cells). Finally, like for pyramidal cells of layers II/III, there were no significant differences in GABAR-PSC and GluR-PSC frequencies in wild-type cells and mutant cells of the layer V at the 3 developmental periods studied ( n = 22 wild type cells/4 animals and n = 19 mutant cells/ 4 animals at PND 28-35, Fig. 9F).

Together these data showed that the variant affected spontaneous synaptic activity only at developmental stages when RGNA was present and particularly in layers II/III. This suggested that GABAergic interneurons contributing to RNGA in these layers may express M current carried by Kv7 channels containing Kv7.2 subunit.

### *KCNQ2^WT/T274M^* mice displayed spontaneous epileptic seizures preferentially at post-weaning than at juvenile stage

In a previous study (Milh et al., 2020), EEG monitoring following surgical implantation of electrocorticographic electrodes demonstrated the presence of seizures in ∼20 % of *KCNQ2^WT/T274M^* mice (mean age PND 33). Based on our current electrophysiological results and given that seizures in patients disappear with age (Boets et al., 2021), we asked whether the occurrence of seizures would be developmentally regulated and higher in post-weaning than in juvenile mice. For this purpose intracranial video-EEG recording were performed in 32 *KCNQ2^WT/WT^* mice (16 at PND18-21 and 16 at PND35-50) and 32 *KCNQ2^WT/T274M^* mice (16 at PND 18-21 and 16 at PND35-50) for a maximum of 48 hrs (see material and methods).

None of the 32 *KCNQ2^WT/WT^* mice (from PND18 to PND 50) recorded showed abnormal behavioral and electrographic activity (Fig.10A). In contrast spontaneous seizures were detected in 60% of *KCNQ2^WT/T274M^* post-weaning mice (10 out of 16 KI mice), all occurring during sleep. Eight mice presented generalized seizures (Fig. 10Ba) with behavioral manifestations. These seizures were often violent, characterized by a two-stage pattern: an initial ∼5s active running phase where mice jumped and ran followed by a 10-20s tonic phase where animals would fall and adopt a tonic posture with limb extensions (see supplemental video 1). The active running phase was accompanied by a 3-5s depolarization shift that saturated the amplifiers. During the tonic phase, a 3-5Hz spike and wave discharge pattern appeared, progressively increasing in amplitude and frequency. At the end of the tonic phase, spike and wave discharge decreased in frequency leaving place to a flattening of the EEG and to the death of 6 mice. Two mice survived such seizures, the EEG activity progressively resuming after 10-20s of flattening. Partial seizures were observed in two mice. In one mouse the seizure consisted of a 3Hz spike and wave activity that was restricted to the right hemisphere (Fig.10Bb). In the other, 5Hz spike and wave activity was only observed in the right hippocampus (fig. 10Bc, supplemental video 2). In both cases, there was no detectable behavioral activity. As described in a previous study (Milh et al. 2020), spikes are observed in wild type and knock-in animals. Spike frequency was quantified in 10 *KCNQ2^WT/T2474M^* mice (6 that survived after seizures plus 4 that did not have seizures during the 48h of recordings) and in 14 *KCNQ2^WT/WT^* mice. Neocortical spikes were detected in 8 *KCNQ2^WT/T274M^* mice (mean rate of 2.98 ± 1.3 spikes/hr; range 0.1-11.7) and hippocampal spikes (mean rate of 3.0 ± 0.9 spikes/hr; range 0.04-8.13) in 9 animals. Spikes tended to occur in clusters, (10-30 spikes per hour during 30mn to 3h), separated by prolonged periods (hours) of quiescence. They were generally detected simultaneously but at different amplitudes on all electrodes (Fig. 10Ca) but some focal spikes could also occur sporadically (fig. 10Cb). In wild-type mice, rare spikes were detected in 6 out of 16 mice, 6 animals having neocortical spikes (mean rate of 1.1 ± 0.5 spikes /hour, range 0.03-3.1) and 3 mice having hippocampal spikes (mean rate of 2. 8 ± 2.5 spikes/hr, range 0.08-7.9).

**Figure 10:**
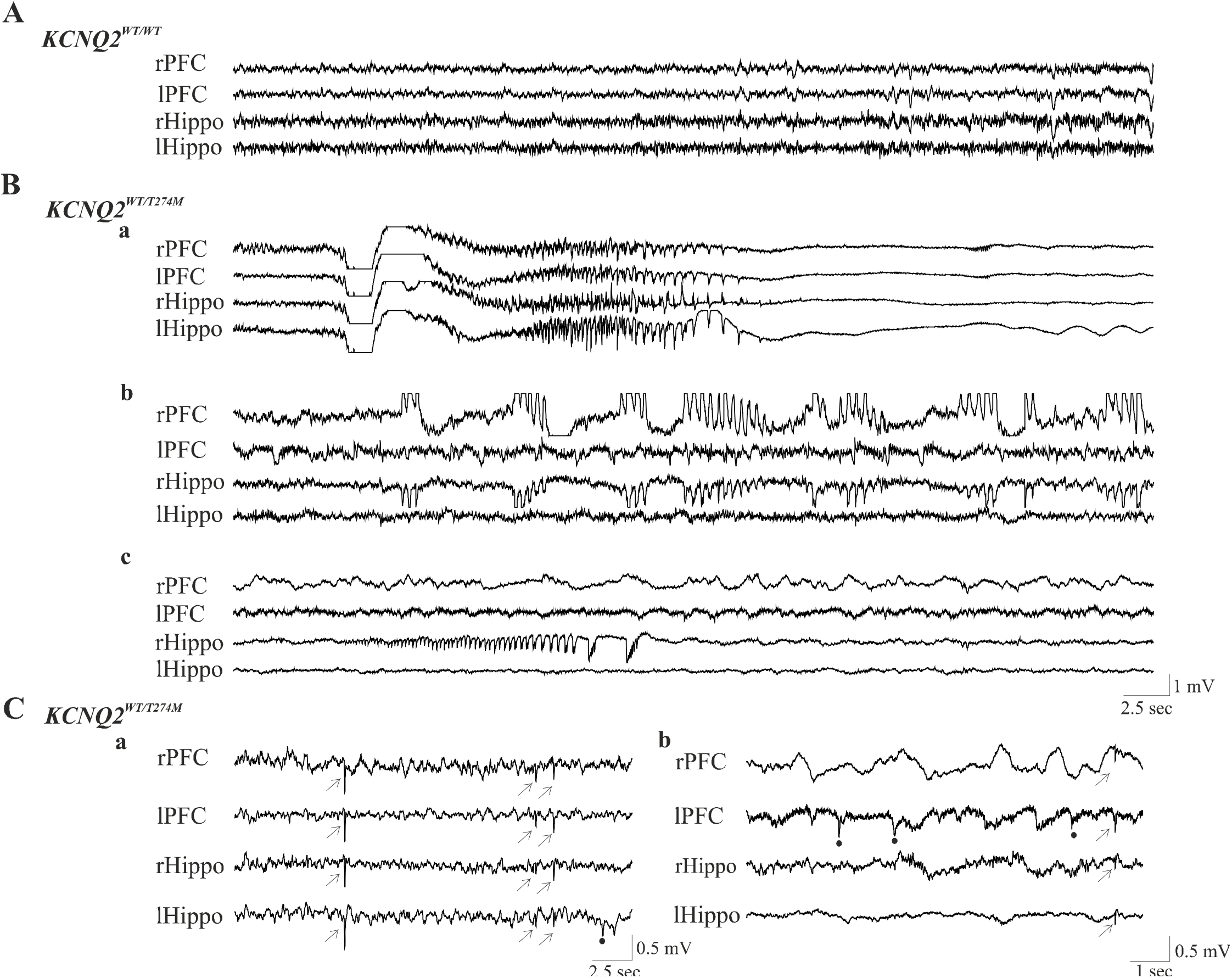
Intracranial EEG –recordings of post-weaning mice Electrodes were implanted in the prefrontal cortex (PFC) and hippocampi (Hippo) of the right (r) and left (l) hemispheres. **A)** Recordings from a *KCNQ2^WT/WT^* mouse. **B)** Spontaneous seizures recorded in *KCNQ2^WT/T274M^* mice. Generalized seizure (**a**). It represented the most often type of seizures encountered in mutant mice. Partial seizures (**b, c**) recorded in two mutant mice with epileptic activities observed in both prefontal cortex and hippocampus of the right hemisphere (**b**) and with an epileptic discharge present only the hippocampus of the right hemisphere (**c**). **C)** Spikes recorded in two mutant mice. These spikes could be synchronized in the right and the left hemisphere (arrows) or be observed in only one structure of one hemisphere (black dot)

At PND 35-50, spontaneous epileptic seizures were recorded in only 25 % of the *KCNQ2^WT/T2474M^* mice (4 out of 16 KI mice). As for post-weaning mice, all seizures were violent, leading to the death of all 4 animals. Spike frequency was also quantified in 12 wild type and 7 knock-in mice. In the neocortex, there were observed in 2 wild type mice (0.11/hr and 2.63 hr) and in 3 knock-in mice at a rate of 4.5 ± 1.3 (range 2.5 -7). In the hippocampus there were detected in 3 wild type mice at a rate of 0.04 ± 0.015/hr (range 0.01-0.06) and in 2 knock-in mice (0.09/hr and 0.11/hr).

Together these data confirmed that the p.T274M variant led to epileptic seizures in mice and indicated that brain excitability of post-weaning KI mice was higher and more prone to generate epileptic seizures than that of juvenile mice.

## Discussion

In this study we performed electrophysiological recordings in a mouse model of *KCNQ2*-related DEE carrying the loss-of-function p.T274M variant. This variant was identified in several patients (Weckhysen et al., 2012; Milh et al., 2013, 2015; Orhan et al., 2014), therefore offering the opportunity to better understand how dysfunction of Kv7 channel affects brain activity during development. Here, we provide evidence that the p.T274M variant lead to epileptic seizures *in vivo* and at cellular level to a decrease of M current, an increase of pyramidal cell excitability and of network driven synaptic events although with some differences depending of the cortical layers. Importantly, as for seizures, we show that these alterations are time-limited, observed until post-weaning stage and not latter with the full recovery of Kv7/M channels function and the normalization of electrophysiological activity at juvenile stage.

Thus, we observed in layers II/III that there was a strong parallelism between the effect of the variant and of the M channel blocker XE-991 on the electrophysiological properties of pyramidal cells until PND19-21. The responses of the cells to the injection of current were similarly impacted by the variant or the blocker with a general increase in neuronal excitability up to a certain level of current injected. These results are in keeping with the function of Kv7 channels in the control of neuronal excitability and in particular the fact that they play fundamental role to limit neuronal firing and to reduce the frequency of the discharge. Moreover, the current ramp protocol which in principle should reveal better the contribution of Kv7 channels on neuronal response over inactivating ionic channels showed that both mutant cells and XE-991 treated wild type cells discharged at higher frequency for lower amount of current injected and generated larger depolarization than in wild type cells. These results support also the idea that the electrophysiological consequences of the variant on cellular properties is related to a decrease in M current and a concomitant increase in membrane resistance at potentials where M current are normally activated. In keeping with this, we observed that M current density and chord conductance were reduced in mutant cells. We cannot exclude the possibility that other conductances are affected in mutant cells and participated to neuronal hyperexcitability as it was recently shown for the calcium-activated BK and SK channels in human iPSC-derived neurons expressing the *KCNQ2* p.R581Q pathogenic variant (Simkin et al., 2021).

Our data also indicated that Kv7 channels control neuronal excitability in motor cortex already during the neonatal period. This was also described in acute dissociated pyramidal cells and paravalbumin like interneurons of layers II/III of somatosensory cortex, CA1 and CA3 pyramidal cells of the hippocampus, lumbar interneurons of the spinal chord from rodents aged one week (Safuilina et al., 2008; Guan et al., 2011; Marguet et al., 2015; Soh et al., 2018;Verneuil et al., 2021). Thus Kv7/M channels appear as important modulators of firing properties of neurons from several regions already at early developmental stage and even if there are expressed to a much lower level than at juvenile and adult stages they are likely to play fundamental role for proper brain development and for preventing hypersynchronicity of cellular activity (Peters et al., 2005; Marguet et al. 2017; Hou et al., 2021).

Our results also suggest that Kv7.2 subunit may strongly contribute to M current in layers II/III. A recent study performed in the somatosensory cortex, showed that the conditional ablation of *KCNQ2* or the transfection by electroporation of the pathogenic p.I205V variant –related to DEE leads to a hyperexcitability of pyramidal cells of layers II/III, whereas conditional ablation of *KCNQ3* did not have such impact (Niday et al. 2017, but see Hou et al. 2021). In the present study, we cannot exclude the contribution of other subunits to Kv7 channels in these cells and in particular Kv7.5 but if any this should be small regarding the large reduction of M current (∼85%) produced by the blocker at 20µM while homomeric Kv7.5 or heteromeric Kv7.3/Kv7.5 subunits association display low sensitivity to XE-991(Schroeder et al., 2000).

The fact that in layers II/III, XE-991 and the p.T274M variant had equivalent consequences on voltage response to current injection was nevertheless surprising regarding their respective effects on M current; the variant reducing current density by ∼30 -40 %. There are two possible explanations at least: - a more efficient blockade of Kv7 channels by XE-991 in voltage clamp than in current-clamp experiments. Indeed XE-991 operates better when channels are open (Greene et al., 2017) a situation that was favored by the protocol used to study M current than to study neuronal excitability. However, this explanation is difficult to reconcile with the more powerful effects exerted by XE-991 over those produced by the variant in the response of layer V pyramidal cells at PND19-21 to the injection of steps or ramp of current. We rather explained these results by the combination of a high sensitivity of pyramidal cells to Kv7 channels activity with a rapid saturation of voltage response /current relation. A similar sensitivity is certainly present in pyramidal cells of the layer V at PND19-21; alterations produced by the variant on both voltage response and M current are within the same range than those produced in layers II/III. Moreover the responses of wild type cells to the injection of steps and ramp of current were equivalent. The stronger effect produced by XE-991 denotes certainly a capacity of voltage response to current injection that is larger in layer V than in layers II/III.

An important difference between layers II/III and layer V pyramidal cells was the lack of effects of the variant and of XE-991 in layer V at PND7-9. This is likely the consequence of the low expression level of M current in wild type cells of the layer V which was twice lower than in layers II/III. It is possible that M current in layer V is carried by channels that include Kv7.5 subunit rather than Kv7.2, the Kv7.3/Kv7.5 association generating lower current than that carried by the Kv7.2/Kv7.3 association (Jentsch, 2000) while the latter would conduct large part of the current at PND19-21 and already since the neonatal period in layers II/III. It is well known that there are differences in the expression of ion channels and intrinsic properties of cells of a same structure. This has been documented in particular in the superficial and deep layers of CA1 and CA3 pyramidal cells, between CA3 and CA1 pyramidal cells or between CA1 pyramidal cells of dorsal and ventral hippocampus (Tzingounis et al., 2010; Mizuseki et al., 2011; Marissal et al., 2012; Hönigsperger et al., 2015; Cembrowski and Spruston 2019). Thus, for example and regarding the composition of Kv7 channels it has been shown that Kv7.2,3,5 subunits coexist in CA3 pyramidal cells but with a high proportion of Kv7.5 subunit which participate to the mAHP and slow AHP in these cells whereas in the CA1 pyramidal cells the expression of Kv7.5 is low and M current is mainly mediated by Kv7.2/Kv7.3 (Tzingounis et al., 2010).

The analysis of M current showed an apparent reversal potential (Erev) of ∼ −74 mV at post-weaning and juvenile developmental stages and even at a more depolarizing potential at PND7-9, mean value that largely deviates from E_K_ calculated by Nernst equation (∼-94 mV). A similar value of M current reversal potential has been described in hypoglossal motoneurons from inward relaxation analysis (Ghezzi et al., 2017), while in the neocortical pyramidal cells or mossy fibers terminals values were close to the calculated E_K_ (Battefeld et al., 2014; Martinello et al., 2019). This discrepancy is difficult to understand, the simplest explanation could be an imperfect clamp of the membrane when employing strong voltage command particularly when the current is generated at distance from the recording site although the AIS is close to the soma. In studies performed by Battefeld and coll. and Martinello and coll., M current was recorded at axonal node (bleb) and at axon boutons respectively, sites that concentrated Kv7 channels and that are of smaller size than the large soma of pyramidal cells where recording electrodes were positioned in the present study. There are also other possible explanations for the depolarized value of Erev in our conditions: i) a contamination by another conductance responsible of an inward current. However channels generating inward current were blocked in our experimental conditions. This is the case of voltage gated Na^+^, and Ca^2+^ channels as well as HCN channels; ii) a permeability of Kv7 channels to Na^+^. Recent observations demonstrate in heterologous cells a physical interaction between Kv7.2 subunit and SMIT1 which enables the transport of myo-inostol into the cells and the raise of PIP2, a phosphoinositide that plays critical role for Kv7 channels function. The formation of this complex increases Na^+^ permeability, which if present in cortical neurons should deviate reversal potential of M current from E_K_ (Manville et al., 2017); iii) As mentioned by Ghezzi et al. (2017), Nernst equation does not take into account the presence of pump, transporters, and non-permeant anionic molecules in the cytoplasm which may affect spatial distribution and concentration of intracellular K^+^ (see also Rahmati et al., 2021). Further studies are necessary to clarify this point.

An important issue of the present study is that the action of the variant on neocortical pyramidal cells activity is time-limited. Neither the membrane response to current injections nor M current density and chord conductance were significantly different in wild type and mutant cells at PND 28-35. The similar level of excitability of mutant and wild type cells is likely related to the lack of effect of the variant on Kv7 channels rather than to a more limited function of these channels in the regulation of neuronal activity since wild-type cells were still sensitive to XE-991at PND28-35. Our results show that M current in wild type and mutant cells seems evolved differently in layers II/III and V at post-weaning and juvenile stages. In layers II/III, I/V and G/V relations were significantly smaller in wild type cells at PND 28-35 than at PND19-21 while these relations were stable in mutant cells so that current density and chord conductance in these cells approached those of wild-type cells at PND28-35. It is the opposite in layer V where at this developmental stage I/V and G/V relations were stable in wild type cells but increased in mutant cells from PND19-21 to PND28-35. This indicates a different mode of regulation for the expression of M current in wild type cells of the layers II/III and V at these two developmental stages. Moreover these results also suggest that at PND28-35 different mechanisms are engaged in mutant cells of layers II/III and V to bring Kv7/M channels function to a physiological level and abolish the effects of the variant on neuronal excitability.

Our data showed that the variant affected not only the properties of pyramidal cells but may also probably affect the properties of interneurons. This was suggested from the analysis of spontaneous synaptic activity. We observed the presence of spontaneous network driven events that were mediated by GABA receptors (called RGNA), and more often observed in layers II/III than in the layer V. We found that RGNA recorded in layers II/III were highly sensitive to the variant and in particular their frequency and number of cells where they have been recorded which were increased in the knock-in mice. Spontaneous network-driven activities have been described in several developing structures including the hippocampus (Ben-Ari et al., 1989, 2007; Garaschuk et al., 1998), the neocortex (Garaschuk et al., 2000; Allene et al., 2008; Modol et al., 2017) where they were called giant depolarizing potential (GDPs) or early network oscillations (ENOs), the spinal cord (Landmesser and O’Donovan et al., 1984), the retina (Meister et al., 1991). In the cortex these patterns modify the efficacy of developing GABAergic and glutamatergic synapses and are likely to play important role in the wiring of neuronal circuit which alteration could be the cause of epilepsy and developmental delay (Ben-Ari et al., 2007; Griguoli and Cherubini, 2017; Khazipov and Milh 2018). Interestingly, in the hippocampus change in the frequency of GDPs (increase or decrease) has been reported in some animal models of neurodevelopmental disorders (Griguoli and Cherubini, 2017). However RGNA display some differences with GDPs/ENOs. Indeed; i) they were still observed in the presence of glutamate receptors antagonists, suggesting they do not necessitate pyramidal cells activation for their generation; ii) they seems largely or exclusively mediated by GABA receptors while GDPs/ENOs contain a glutamatergic component (Khalilov et al., 2015). The fact that pyramidal cells were silent (in cell –attached configuration) and did not fire action potential during bursts of GABAR-PSCs even at PND 7 when GABA exerts its depolarizing action (Ben-Ari et al. 2007) explain probably why glutamate receptors did not contribute to RGNA; iii) RGNA are observed at PND19-21, in particular in layers II/III and even with an higher frequency than at PND7-9 while GDP/ENOs disappeared in cortical slices from rodents aged one-two weeks. It is possible that the differences in the properties of RGNA and GDPs/ENOs are related to the cortical structure analyzed and/or the rodents species used. In the hippocampus synaptic-driven network patterns are orchestrated by a subpopulation of pioneer early born gabaergic interneurons expressing somatostatin, possessing widespread axonal arborization and acting as hub cells (Bonifazzi et al., 2010; Picardo et al., 2011). Existence of GABAergic hub cells has also been described in the entorhinal cortex and they are likely to exist in other neocortical regions such as the motor cortex to orchestrate network activity (Modol et al., 2017). We speculate that Kv7 channels incorporating Kv7.2 subunits play important role in the firing properties of these cells. However the reasons why the variant impacts RGNA in layers II/III but much less in the layer V is difficult to understand notably because some interneurons including hub cells of both layers may be interconnected which would allow the propagation of RGNA from one layer to another (Jiang et al., 2015; Modol et al., 2017). There are at present some data regarding the expression and function of Kv7 channels in interneurons. Studies have essentially been performed in hippocampal interneurons where expression and modulation of neuronal firing by Kv7 channels have been demonstrated for regular and fast spiking interneurons maintained in primary hippocampal cultures (Grigorov et al., 2014), in a subset of somatostatin oriens-lacunosum moleculare (OLM) interneurons and paravalbumin expressing interneurons in the CA1 region of the hippocampus; in the basket cells, OLM and bistratified interneurons of the CA3 region (Lawrence et al., 2006; Nieto-Gonzalez and Jensen 2013; Fidzinski et al., 2015; Soh et al.2018). Other data provided also evidences for the modulation of neuronal firing by Kv7 channels of paravalbumin expressing interneurons in layers II/III of the somatosensory cortex (Soh et al., 2018). It is therefore not surprising that Kv7.2 variants affect spontaneous gabaergic activity in the neocortex (the present study) and the hippocampus (Uchida et al, 2017, but see Marguet et al., 2015). Although it is unclear if RGNA have any correspondence *in vivo*, our data suggest that a population of interneurons, which identity and properties remain to be characterized, are impacted by the variant and must play important role in the pathophysiology of *KCNQ2*-related DEE.

Video/EEG monitoring revealed the presence of seizures with or without behavioral manifestation. As described in previous studies (Milh et al., 2020), behavioral seizures, which account for the majority of epileptiform events recorded, were associated with violent locomotor activity and mostly terminated with the death of the animal. The synchronous spike-and-wave discharge pattern associated with such seizures is reminiscent of the patterns typically observed in generalized epilepsies. However, we also observed focal seizures, restricted to one hemisphere or one hippocampus. These seizure patterns were not associated with any behavioral manifestations. Despite the presence of seizures, we did not observe the suppression burst patterns that are typically observed in *KCNQ2*-related DEE patient in neonatal period. However, given that our recordings were only performed after weaning (see materials and methods), it is possible that we missed the temporal window where such patterns would occur in mice.

In summary, we showed that the p.T274M variant clearly affected the properties of cortical cells and ongoing synaptic activity which may contribute to epileptic seizures. Our data also show that the electrophysiological consequences of Kv7 channels dysfunction *in vivo* and *ex-vivo* is transient and observed during a limited period of development. These findings are particularly relevant to *KCNQ2*-related DEE knowing that for a majority of patients a normalization of the electrographic activity and remission of the epilepsy few weeks to several months after seizures onset are often observed while neurological outcome remains poor. In particular the epilepsy of one patient carrying the p.T274M variant was active for only 6 weeks, controlled by anti-epileptic drugs then in remission without treatment from the age of 1 year (Milh et al., 2013, see also Wechuysen et al., 2012). It is tempting to speculate that the normalization of the EEG also results from recovery of Kv7/M channels function. However in spite of this, the patient made almost no motor acquisition and displayed intellectual disability (Wechuysen et al., 2012; Milh et al., 2013). Interestingly, *KCNQ^WT/T274^* mice display important cognitive deficits that persist in adulthood (Milh et al., 2020) and we observed that seizures were preferentially recorded in post-weaning mice and much less in juvenile mice. Although several questions remain regarding the role of neonatal seizures and early patterns of cortical activity in brain development (Khazipov and Milh, 2018; Bitzenhofer et al., 2021), the elucidation of mechanisms of Kv7/M channels recovery and reasons why this occurs at a particular stage of development represent important aspect of future studies for the understanding of the pathophysiology of *KCNQ2*-related DEE. In addition, this may have implication to develop therapies with adapted target and to determine the most appropriate time window at which they should be installed to prevent long-term neurological deficits.

## Materials and Methods

### Animals and genotyping

All experiments were carried out in accordance with the European Communities Council Directive of September 22, 2010 (2010/63/UE) related to laboratory animals used for research purposes. The study was approved by ethics commity of the “Ministère de l’Enseignement Supérieur de la Recherche et de l’Innovation (APAFIS#27891-2020110514571312 v2). Experiments were performed on male and female postnatal day 7 (PND 7) to post-natal day 50 old 129Sv mice, housed in a temperature-controlled environment with a 12 light/dark cycle and free access to food and water (INMED animal facilities).

The generation of heterozygous *KCNQ2^WT/T274M^* knock-in mice and genotyping protocols have been described in a previous paper by Milh et al. (2020). Briefly, mice were genotyped by PCR. The following two primers were included in the PCR: A common forward primer (5’-CTTGATCTTGTCCCCTGACTTGGTAGG-3’) and a common reverse primer (5’-CCTAACATCTCCAGAGTAGGAAGGTGCG-3’). The primers amplified a 603 bp wild-type fragment and/or a 688 bp mutant fragment.

### Slices preparation

Experiments were performed on pyramidal neurons located in layers II/III and V of motor cortical slices.

Mice were euthanized by decapitation. The brain was rapidly removed and placed in an oxygenated ice-cold choline solution containing (in mM): 132.5 choline chloride, 2.5 KCl, 0.7 CaCl_2_, 3 MgCl_2_, 1.2 NaH_2_PO_4_, 25 NaHCO_3_ and 8 glucose; oxygenated with 95% O_2_ and 5% of CO_2_. Coronal slices (300 μm-thick) were cut using a vibratome (Leica VT1200S; Leica Microsystems, Germany) in ice-cold choline solution oxygenated with 95% O_2_ and 5% of CO_2_. Before recording, slices were incubated in an artificial cerebrospinal fluid (ACSF) solution with the following composition (in mM): 125 NaCl, 3.5 KCl, 2 CaCl_2_, 1.3 MgCl_2_, 1.25 NaH_2_PO_4_, 26 NaHCO_3_ and 10 glucose equilibrated at pH 7.3-7.4 with 95% O_2_ and 5% CO_2_ at 34 °C for 20 min and then at room temperature (22–25 recovery. Slices were placed into the recording chamber for electrophysiological purpose where they were fully submerged and superfused with oxygenated ACSF solution at 34- °C at a rate of 5 ml/min.

### *Ex-vivo* electrophysiology

Pyramidal cells of layers II/III and V were recorded under visual control with a Zeiss Axioscope 2FS microscope in cell-attached and whole cell configurations using borosilicate glass capillaries (GC 150F-15). For recordings in current-clamp mode, patch-pipettes were filled with a solution containing (in mM): 140 KMeSO_4_, 6 NaCl, 10 HEPES, 1 MgCl_2_, 4 Mg-ATP, 0.4 Na_2_-GTP. The pH was adjusted to 7.35 with KOH. The resistance of the pipettes was of 5-6 MΩ. For recordings of spontaneous synaptic activities in voltage-clamp mode, patch pipettes were filled with a solution containing (in mM): 120 Cs-gluconate, 10 CsCl, 10 HEPES, 4 Mg-ATP, 0.4 Na_2_-GTP. The pH was adjusted to pH 7.35 with CsOH. The resistance of the pipettes was of 6-7 MΩ. For recordings of M current, patch pipettes were filled with a solution containing (in mM): 135 K-gluconate, 10 KCl, 10 HEPES, 5 phosphocreatine, 4 Mg-ATP, 0.4 Na_2_-GTP. The pH was adjusted to pH 7.35 with KOH. The resistance of the pipettes was of 5-7 MΩ.

All reported potential values were corrected for the liquid junction potential, calculated to be ∼2 mV with the KMeSO_4_ pipette solution and ∼15 mV with the Cs-gluconate and K-gluconate filled pipette solution.

Recordings in current-clamp mode were performed in presence of 2,3-Dioxo-6-nitro-1,2,3,4-tetrahydrobenzo[*f*]quinoxaline-7-sulfonamide (NBQX, 10 µM) to block AMPA/Kainate receptors; D-2-amino-5-phosphonovalerate (D-APV, 40 µM) to block NMDA receptors; 6-Imino-3-(4-methoxyphenyl)-1(6*H*)-pyridazinebutanoic acid hydrobromide (SR 95531, Gabazine 5 µM) to block GABA_A_ receptors. Some experiments were performed in continuous presence of Kv7 channels blocker 10,10-*bis*(4-Pyridinylmethyl)-9(10*H*)-anthracenone dihydrochloride (XE-991, 20µM). In these slices, pyramidal cells recordings were performed ∼20-30 min after starting the superfusion with XE-991 (see Hu and Bean, 2018).

Resting membrane potential was determined in current clamp as the potential upon break-in, before any current injection. Action potential (AP) threshold was defined as the membrane potential at which the rate of depolarization is maximal. All measurements were filtered at 3 KHz using an EPC10 amplifier (HEKA Electronik, Germany) and sampled at 10 kHz. Data were analyzed off-line using Clampfit (Molecular Devices), miniAnalysis (Synaptosoft), Origin 9 (Origin Lab), Prism 6 software (GraphPad).

Only cells with a stable resting membrane potential more negative than −60 mV were used in this study. All measurements in current-clamp mode were performed from a membrane potential of −70 mV (-72 with LJP correction), and if necessary, current was injected during the experiment to keep this value constant.

To calculate the membrane input resistance (Rm), voltage responses to the injection of five depolarizing and hyperpolarizing current steps, with an increment of +/-5 pA and applied during 500 msec, were fit with a linear function to yield the slope value.

To calculate the membrane time constant (τ_m_), e voltage response to the injection of a hyperpolarizing current step of −20 pA for 500 msec was fitted with a single exponential function (Origin 9) to yield the tau value. Membrane capacitance (Cm) was then calculated according to the equation C_m_ = τ_m_/R_m_.

Action potentials were elicited by injection of short 10 msec and long 1 sec depolarizing current steps of 20 to 300 pA (in 20 pA increments) in pyramidal cells from animal aged one week and of 50-750 pA (in 50 pA increments) from older animals. Currents injected by a slow ramp protocol were of 300 pA and 500 pA applied in 10 sec in pyramidal cells from animal aged one week and from older animals respectively.

M current was recorded in presence of : NBQX (10 µM), D-APV (40 µM), SR 95531 (5 µM), tetrodotoxin (TTX, 1µM) to block voltage gated Na^+^ channels, CsCl (2 mM) to block HCN/ I_H_ channels, 4-aminopyridine (4-AP, 2 mM) to block Kv1/ I_D_ channels and Kv4/I_A_ channels and CdCl (200 µM) to block voltage-gated Ca^2+^ channels and calcium dependent potassium channels.

In order to open Kv7 channels, a slow ramp of voltage was applied from −90 mV to reach +10 mV in 30 sec. Cells were maintained for 1 sec at this membrane potential and 1 sec hyperpolarizing voltage steps down to −90 (in 10 mV increments) were applied to deactivate the channels. A short (100 msec) hyperpolarizing voltage steps command of 10 mV was applied at −90 mV before each slow ramp of voltage to ensure the stability of the access resistance and the quality of the recordings (Supplemental Fig. 3A). M current amplitude was measured as the difference between the instantaneous (Iins) and steady-state current (Iss) levels at the onset and at the end of each hyperpolarizing voltage steps applied from +10 mV respectively (supplemental Fig. 4D). The chord conductance (G) values were obtained from the deactivated current amplitude divided by the driving force for K^+^ (G= I/V-E_K_) where V is the membrane potential at which the current was measured, E_K_ is the reversal potential of K^+^ ions calculated by Nernst equation ( ∼-94 mV in our recording conditions).

To measure the current deactivation kinetics, current traces were fitted with a single or a double exponential function of the following form: y=Afastexp(τ / τfast)+Aslowexp(τ / τslow) (where Afast and Aslow are the fractions of the fast and slow component of the current and τfast and τslow are the respective fast and slow time constant). The time constant representing the weighted average of the fast and slow components of current deactivation was calculated with the following equation τ slowAslow)/ (Afast +Aslow). Currents were analyzed using Origin 9.0 software.

### *In vivo* Video-EEG monitoring

For ethical reasons we performed video-EEG monitoring for a maximum of 48 h after weaning. Tethered recordings would require isolating the pups from the dam before weaning and radio-telemetry devices are too large for implantation in animals younger than PND 60. For these reasons, recordings were performed starting from post weaning age (PND18-20) and using custom made tether-preamplifier assemblies allowing mice to move freely. The recording apparatus was adapted by the manufacturer (Neuralynx, Bozeman, MT). It consisted of a 64-channel Digitalynx SX (24 bit A/D converter) connected to a 4-way cable-splitter. Each input cable was connected to a 16-channel headstage pre-amplifier that allowed to record signals from up to four freely moving mice at a time. An ethernet camera (Basler acA1300-75gc) was placed above the cages and connected to a Gigabit ethernet acquisition card (Neuralynx, Bozeman, MT). The recording software interface (Cheetah 6.4.1) allowed to simultaneously record intracranial local field potentials (LFPs) from all mice synchronously with the video signal from the camera. LFPs (recorded against the ground) and video data were acquired at 1 KHz and 30 Hz, respectively.

Electrode arrays consisted of four 50µm nichrome, formvar insulated wires bent and cut to the desired lengths. A 76µm stainless steel wire (A-M system) serving as a ground was bent perpendicularly and cut at 1.5mm. During surgery, the tip of this wire was de-insulated for 1mm. All wires were connected to a board-to-board connector (Mill-Max Manufacturing Corp).

### Surgery

Thirty minutes before surgery, mice were anesthetized with 4% isoflurane and injected with buprenorphine subcutaneously (Buprecare, 0.03 mg/kg). They were then re-anesthetized (4% isoflurane for induction decreased progressively to 1.5 % for anaesthesia maintenance), placed in the stereotaxic apparatus (Kopf Instruments) and breathing through an anaesthesia mask (TEM Sega, France). The scalp was shaved and cleaned with three successive treatments of betadine and 70 % ethanol. Body temperature was maintained at 36 °C through the whole procedure with a heating pad placed under the animal and monitored with a rectal probe (FHC). An ophthalmic gel was placed on the eyes (Lanolin and Retinol mix, Vitamin A Dulcis, France) to prevent dryness. Lidocaine 5 % was injected subcutaneously under the scalp, 5 min before incision. The scalp was then removed and the bone surface cleaned. Small incisions were performed on the bone surface and holes were drilled at the following coordinates: at PND 18-20: Prefrontal cortex (PFC): AP= +0.1; L=+/− 0.09; DV=1.5; Hippocampus : AP= −0.23; L=+/− 0.25; DV=2.5; at PND34-50. PFC: AP=+0.12; L=+/−0.12; DV=1.5; Hippocampus: AP=-0.23; L=+/− 0.27; DV=2.5mm with regards to bregma – AP=anteroposterior, DV=dorso-ventral; L= Lateral) with regards to bregma – AP=anteroposterior, DV=dorso-ventral; L= Lateral).

Electrode wires were bent 90° 1mm before the tip and inserted through the craniotomies. The ground wire was placed above the cerebellum, through a hole in the dorsal skull suture. Veterinary grade cyanoacrylate gel (Vetbond, 3M) was used to secure electrodes on the skull and spread throughout the skull. Dental cement (Paladur, Kulzer) was then used to secure the body of the implant to the skull.

### Video-EEG monitoring

On the day of recordings, up to four weaning (P18-20) or adult (P34-50) mice were connected to the cable-headstage assembly. They were allowed to move freely in their home cage (one animal per cage, 33,2×15×13 cm) and had access to standard food diet and non-wetting water gel for hydration (Hydrogel-ClearH2o, Westbrook ME, USA). Recordings were started in the afternoon following surgery. Animals were inspected regularly during the day (every 2h) to ensure they did not have any adverse reactions to the surgery and recording procedure. To maintain recordings to a reasonable size, a Matlab (Mathworks, MA) routine was used to remotely control the Cheetah acquisition software (Neuralynx, Bozeman, MT) and start a new recording every half hour.

Mice were euthanized at the end of the 48h of video-EEG recordings using isoflurane anaesthesia (5 %) follow by cervical dislocation. The brain was then removed and cut to verify the position of the electrodes.

### Data analysis

It was performed using custom made Matlab routines and graphical user interfaces. LFP signal was bandpass filtered (1-400Hz) and putative interictal spikes (IIS) were automatically detected using White et al., (2006) method. Briefly an IIS was considered each time the slope of signal over 16ms was greater than 6 times the standard deviation and its amplitude greater than 3 times the standard deviation of the local field potential. Then, putative IIS were then visualized together with the corresponding video. Scratching, chewing or bumping artefacts were therefore detected and removed from analysis. Other types of electrical artefacts, characterized by vertical slopes and symmetrical shapes, were also discarded. Seizures were considered when more than 20 IIS were detected in a 10s window. <u>Statistics</u>

Data are represented as means ± S.E.M. Student’s t-test was used to compare means of two groups when the data’s distribution was normal. When the normality test failed, we used the non-parametric Mann– Whitney test for two independent samples. The normality was assessed using Shapiro-Wilk test. Two-way Anova test was used to assess the statistical significance of two graphs. Statistical analysis was performed using Graphpad Prism software. ns: not significant; *^,§^ p < 0.05; **, ^§§^p <0.01; and ***^,§§§^ p < 0.001.

## Acknowledgements

This work was supported by INSERM (Institut National de la Santé et de la Recherche Médicale), by the Agence National pour la Recherche (ANR −19-CE17-0018-02, IMprove), by the Ministère de l’Enseignement Supérieur, de la Recherche et de l’Innovation (NB). We would like to thank Aurélie Montheil and Francesca Bader for mice genotyping (PBMC at INMED) and INMED’s animal facility technicians for excellent technical support. We thank also Dr Jerôme Epsztein for his constructive remarks on the manuscript.

---

**<u>Supplemental Video 1</u>:** Video-EEG monitoring of a generalized seizure in a KI mice

**<u>Supplemental Video 2:</u>** Video-EEG monitoring of a partial seizure with spike and wave activities in the right hippocampus and without visible behavioral manifestation.

